# An integrated picture of the structural pathways controlling the heart performance

**DOI:** 10.1101/2024.05.11.593706

**Authors:** Ilaria Morotti, Marco Caremani, Matteo Marcello, Irene Pertici, Pasquale Bianco, Theyencheri Narayanan, Gabriella Piazzesi, Massimo Reconditi, Vincenzo Lombardi, Marco Linari

**Affiliations:** PhysioLab, University of Florence, 50019 Sesto Fiorentino, Italy; Department of Biology, University of Florence, 50019 Sesto Fiorentino, Italy; European Synchrotron Radiation Facility - The European Synchrotron, Grenoble 38043, France; Department of Experimental and Clinical Medicine, University of Florence, 50134 Florence, Italy

## Abstract

Regulation of heart function is attributed to a dual filament mechanism: (*i*) the Ca^2+^-dependent structural changes in the regulatory proteins of the thin, actin-containing filament making actin available for myosin motor attachment^1^, and (*ii*) the release of motors from their folded (OFF) state on the surface of the thick filament^2^ allowing them to attach and pull the actin filament. Thick filament mechanosensing is thought to control the number of motors switching ON in relation to the systolic performance^3^, but its molecular basis is still unknown. Here high spatial resolution X-ray diffraction data from electrically paced rat trabeculae and papillary muscles call for a revision of the mechanosensing hypothesis and provide a molecular explanation of the modulation of heart performance also in light of the recent cryo-EM thick filament structure^4, 5^. We find that upon stimulation titin activation^6^ triggers structural changes in the thick filament that switch motors ON throughout the filament within ∼½ the maximum systolic force. These structural changes also drive MyBP-C N-terminus to bind actin^7^ promoting first motor attachments from the central 1/3 of the half-thick filament. Progression of attachments towards the periphery of half-thick filament with increase in systolic force is carried on by near-neighbour cooperative thin filament activation by attached motors^8^. The identification of the roles of MyBP-C, titin, thin and thick filaments in heart regulation enables their targeting for potential therapeutic interventions.

## INTRODUCTION

Muscle contraction is the generation of force and shortening due to cyclic ATP-driven interactions between the motor protein myosin II and actin. At the level of the sarcomere, the structural unit of the striated (skeletal and cardiac) muscle (Fig. 1a), myosin motors (orange) extending in two bipolar arrays from the thick filament (light blue) at the centre of the sarcomere (M-line) bind the actin monomers on the overlapping thin filaments (yellow), originating from the Z-line bounding the sarcomere. Actin-attached motors undergo the working stroke that induces force development and filament sliding. The contraction is regulated by mechanisms on both thin and thick filaments. In the thin filament, Ca^2+^-dependent structural changes of the regulatory troponin complex (Tn, Fig. 1b light and dark gray and dark brown) and tropomyosin (Tm, red) make the actin sites available for binding the myosin motors shortly after the action potential^1^. On the other hand, in the muscle at rest myosin motors lie on the surface of the filament folded on their tails (blue, Fig. 1b) in the interacting heads motif (IHM^2, 9^), unable to bind actin and hydrolyse ATP^10^.

**Fig. 1.**
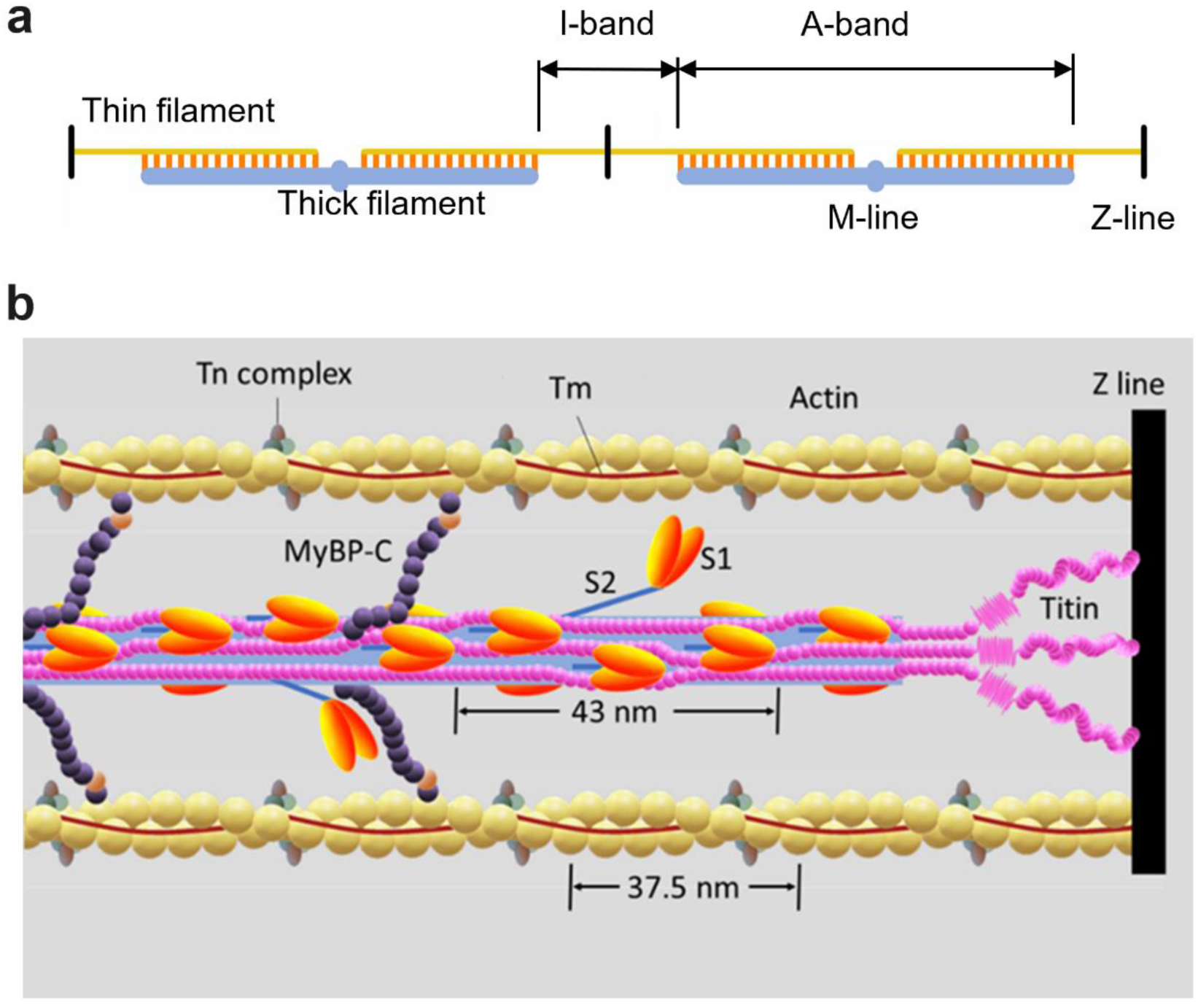
Overview of the sarcomere with its protein content. **a**. Schematic representation of the sarcomere at full overlap (∼2.3 μm sarcomere length) between the thin filament (yellow) and the array of myosin motors (orange) on the thick filament (light blue). **b**. Portion of a half-sarcomere showing the protein assembly. On the thin filament: the two-stranded actin filament with ∼37.5 nm half-helical periodicity (yellow) and the regulatory proteins tropomyosin (Tm, red) and troponin complex (Tn, light and dark gray and dark brown). On the thick filament (light blue): S1 globular head domains of myosin dimer (orange) lying, at rest, on three stranded helical tracks with ∼43 nm periodicity folded back on their S2 tail domains (blue) in the OFF state. Upon activation heads move away with tilting of their tails (ON state, two dimers shown); MyBP-C (dark violet) lying on the thick filament with the C-terminus and extending to the thin filament with the N-terminus, where the phosphorylatable M-domain is brown; titin (magenta) connecting the Z line (black) to the tip of the thick filament in the I-band and running on the surface of the thick filament up to the M-line in the A-band. Only three of the six titin molecules and two of the three series of MyBP-C molecules with 43 nm axial periodicity are represented. The distal portion of the thick filament without MyBP-C has been shortened.

By using X-ray diffraction on intact preparations from skeletal and cardiac muscles it has been shown that during contraction the motors move away from the surface of the thick filament and become available for interaction with actin in relation to the load of the contraction, suggesting a thick filament mechanosensing-based regulation^3, 11–13^. However, growing evidence suggests that Ca^2+^ too can play a role in thick filament activation, even if it is not clear the mechanism and whether the effect is direct^14^ or mediated by accessory and cytoskeleton sarcomeric proteins, like Myosin Binding Protein-C (MyBP-C, Fig. 1b, dark violet) and titin (magenta)^6, 15, 16^. MyBP-C lies on the central 1/3 of each half-thick filament (C-zone) and with its N-terminus can bind either the motor at the head-tail junction on the thick filament or actin on the thin filament establishing an interfilament communication path regulated also by the degree of phosphorylation of its M domain (Fig 1b, brown)^17–21^. Titin spans the whole half-sarcomere, connecting the Z-line with the tip of the thick filament in the I-band, and running associated to the thick filament up to the M-line at the centre of the sarcomere in the A-band^22–24^. Both MyBP-C molecules and A-band titin have fundamental ∼43 nm axial periodicities that match the helical periodicity of myosin motors on the surface of the thick filament (Fig. 1b; ^4, 5, 25, 26^) providing intermolecular interactions with the myosin tail and IHM that influence the regulatory state of the thick filament. Notably, deteriorations of thick filament regulation related to mutations in these sarcomeric proteins and in the myosin itself are common causes of myopathies^27^, and therefore detailed understanding of their function is a fundamental prerequisite for implementation of therapeutic interventions.

In the heart the mechanical activity (systole) consists of short periodic contractions (twitches) triggered by single action potentials and ensuing transitory intracellular calcium rise, during which the blood is pumped by ventricles into the arterial circulation. In the resting period between two systoles (diastole), the heart is filled by the blood from the venous return. The calcium transient following the action potential does not reach the level for full thin filament activation, so that the mechanical response depends from one side on both the internal [Ca^2+^] and the Ca^2+^ sensitivity of the filament^28, 29^, and from the other on the cooperative action of attached motors on thin filament activation^8, 30^. The twitch force can be modulated (inotropic effect) through several regulatory systems, either extrinsic, like neuro-humoral control of the degree of phosphorylation of the regulatory and cytoskeleton proteins as the regulatory light chain in the myosin itself ^31^, MyBP-C ^18, 20, 21, 32, 33^ and titin^34^, or intrinsic, like the sarcomere length (SL) (a property known as length-dependent activation [LDA]^35, 36^). On the other hand, in intact trabeculae the OFF state of the myosin filament in diastole^37^ was perturbed neither by the degree of phosphorylation of sarcomeric proteins nor by the increase in SL, in agreement with the concept of thick filament mechanosensing acting downstream with respect to the diastole^3, 12^.

Regulation of systolic performance based on mechanosensing requires at least two conditions to exist: a fraction of constitutively ON motors in the distal region of the thick filament that triggers the mechanism^11, 38^ and a linear relation between force and degree of thick filament activation^38^. Instead, it is not yet known if and how the structural heterogeneity of the half-thick filament, in which the central 1/3 (C-zone) contains the MyBP-C, integrates with the regional hierarchy of thick filament activation dictated by mechanosensing^3, 13^. Moreover, both X-ray diffraction^12^ and florescent probes^38^ show that motor recruitment from the OFF state attains its maximum earlier than force.

Here we use high spatial resolution X-ray diffraction from intact trabeculae and papillary muscles of the rat ventricle and the disordering effect of lowering temperature^39, 40^ to determine the regional difference in the regulatory state of the thick filament in diastole. Using this approach, we can check if and how the regional variation in the degree of thick filament activation reflects on the axial distribution of motors responsible for the systolic force. At 27 °C, a temperature at which the OFF state in diastole is fully preserved only in the C-zone, the structural signals show that thick filament activation progresses starting from the periphery of the filament and rapidly spreads along the filament for systolic forces ≤1/2 the maximum force. This switching ON of motors throughout the thick filament is preceded by titin-related X-ray signals suggesting that the driving force that perturbs the folded OFF disposition of motors is the Ca^2+^-dependent activation of titin^6^. At the lowest forces motor attachment to actin is limited to the C-zone that is the region with the preserved OFF state in diastole. This calls for a specific action of the N-terminus of MyBP-C on the regulated thin filament^7, 41–43^ promoting first attachments of the myosin motors. At the subsaturating level of intracellular [Ca^2+^] of cardiac myocytes, the firstly attached motors in the C-zone trigger the near-neighbour cooperative activation^8, 44^ that makes actin monomers available for further motor attachment from the progressively more peripheral zones at higher systolic forces.

## RESULTS

### Regional hierarchy of motors disordering by cooling

Lowering temperature below the physiological level is known to induce disorder in the OFF state of the thick filament. Here it is used as a tool to establish a regional hierarchy in the regulatory state of the filament at rest, as elucidated by small angle X-ray diffraction from intact trabeculae and papillary muscles of the rat ventricle (Extended Data Fig. 1a). Lowering the temperature from 35 to 10 °C is accompanied by an overall reduction of the intensities of all reflections.

The intensity of the equatorial 1,0 reflection (*I*_1,0_, Extended Data Fig. 1b, filled circles), associated with the lattice planes containing the thick filaments, decreases when lowering the temperature below 27 °C, and at 10 °C it is halved with respect to the 27-35 °C value. The intensity of the 1,1 reflection (*I*_1,1_, Extended Data Fig. 1b, open circles), associated with the lattice planes containing thick and thin filaments, decreases significantly (by ∼20%) only by cooling to 10 °C. The larger decrease of *I*_1,0_ with respect to *I*_1,1_ is responsible for the increase of the ratio *I*_1,1_/*I*_1,0_ with cooling below 27 °C (from 0.31 at 27-35 °C to 0.47 at 10 °C, Extended Data Fig. 1c), signalling myosin motor movement away from the surface of the thick filament towards the thin filament^45^. Lowering temperature has little effect on the inter-filamentary spacing, measured as the separation of the 1,0 lattice planes, that remains between 35 and 36 nm (Extended Data Fig 1d).

The first order myosin-based layer line (ML1), originating from the myosin motors lying on the surface of the thick filament in three-stranded helical tracks with periodicity 43 nm^46^, shows a monotonic decrease in intensity with cooling (Fig. 2a and d, see also ^40, 47^) indicating a progressive loss of the helically ordered disposition of myosin motors along the filament following their movement away from the filament surface.

**Figure 2.**
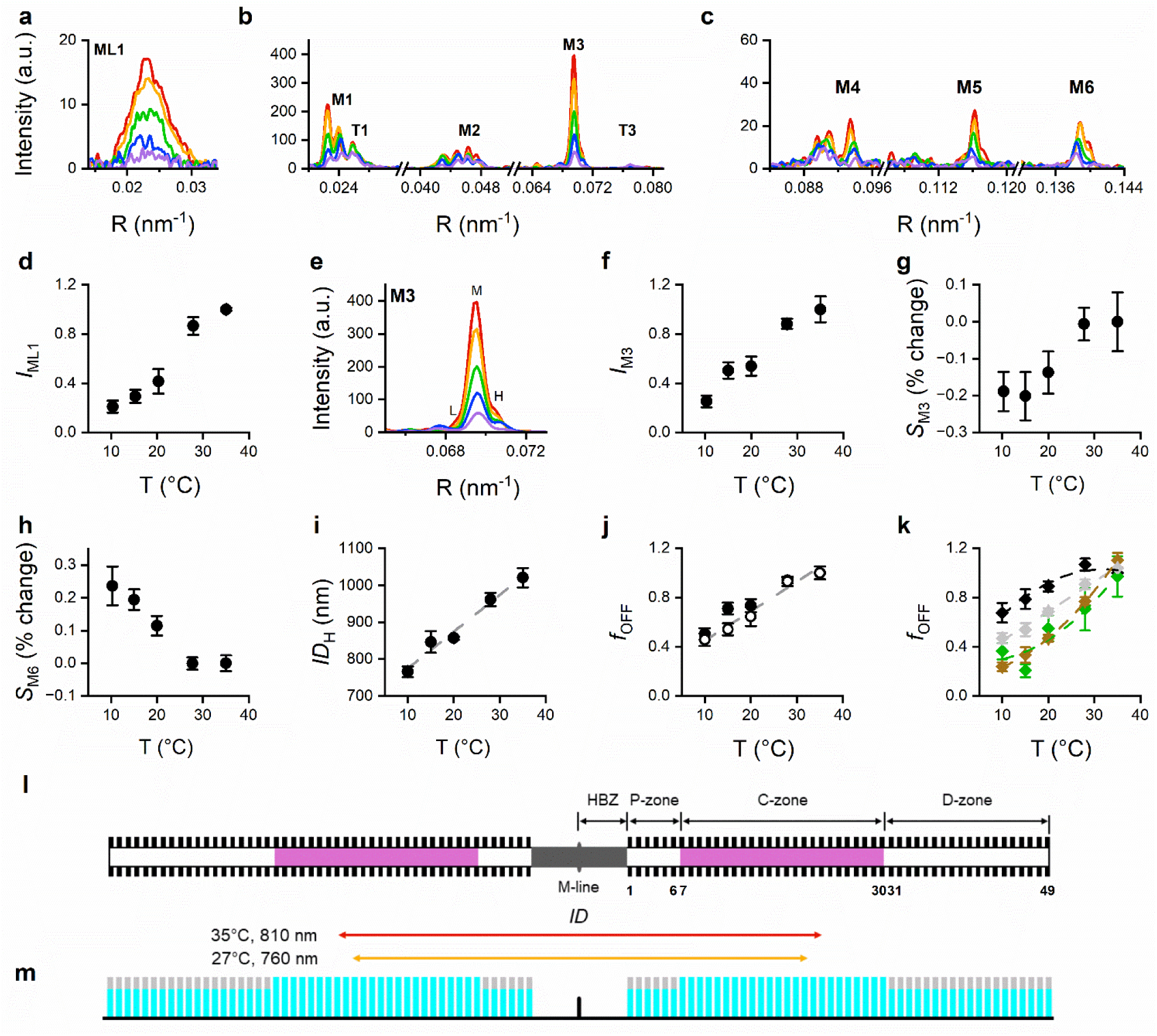
X-ray signals showing the effects of cooling on the structure of thick filament at rest. **a.** Superimposed intensity distribution on the reciprocal space (R) of the ML1 layer line at different temperatures (red, 35 °C; orange, 27 °C; green, 20 °C; blue, 15 °C; violet, 10 °C). **b, c.** Superimposed intensity distributions of the meridional reflections (same colour code as in a). The M (myosin) and T (troponin) series reflections are indicated. **d.** *I*_ML1_, relative to the value at 35°C **e.** Enlargement of the M3 profiles in b (same colour code as in a). L, M and H low, medium and high angle peaks. **f**. *I*_M3_. **g, h.** *S*_M3_ (g) and *S*_M6_ (h) as percentage change relative to the value at 35 °C. **i.** *ID*_H_ measured as the reciprocal of the separation between M and H peaks. The gray line is a linear fit on the data for guiding the eye. In a-c and e each trace is the sum of two exposures 10 ms each. a.u., arbitrary units. In d and f the intensities are normalised for their value at 35 °C. Mean ± SEM, n = 3-7, 3 preparations. **j.** Fraction of motors in the OFF conformation (*f*_OFF_) determined from the square root of either *I*_M3_ (filled circles, from f) or *I*_ML1_ (open circles from d). Values relative to *f*_OFF_ at 35 °C. Mean ± SEM. Gray dashed line, simulated relation from k. **k.** Model prediction of the fractions of OFF motors in the P-, C- and D- zones (*f*_OFF,P_, green, *f*_OFF,C_, black, *f*_OFF,D_, brown, respectively). Gray diamonds, overall fraction of OFF motors. Values relative to the fraction in the C-zone at 35 °C. The dashed lines are parabolic fits on the data (identified by the colour) for guiding the eye. Mean ± SEM, model tested on data from three preparations. **l.** Location on the thick filament of the two bipolar arrays of 49 crowns of motors generating the M3 fine structure. The magenta stripe marks the MyBP-C containing zones from density layer 7 to layer 30; the double-headed arrows below indicate *ID* of the two arrays of diffractors estimated by the M3 fine structure at 35 °C (red) and at 27°C (orange). **m**. Schematic representation of the probability of each layer to be in either the OFF (*f*_OFF_, cyan) or the ON state (*f*_ON_, gray) at 27 °C, represented by the length of the corresponding colour in each bar.

The myosin-based meridional reflections (M1-M6) are orders of a fundamental axial periodicity of ca 43 nm associated to the quasi-regular repeat along the thick filament of density layers due to the crowns of myosin motors and are split in closely spaced peaks originating from the X-ray interference between the two arrays of myosin motors in each thick filament (Fig. 2b, c; ^48^). Among them, the strong M3, originating from the axial repeat of myosin motors with periodicity 14.35 nm (Fig. 2b,e, ^3^), reduces its intensity (*I*_M3_) with cooling throughout the whole range of temperatures, and *I*_M3_ at 10 °C is 1/4 its value at 35 °C (Fig. 2f). This reduction is similar to that of *I*_ML1_, indicating that together with the loss of the helically ordered disposition of motors on the surface of the filament, there is a reduction of their axially ordered folded state. The fine structure of the M3 reflection (Fig. 2e) at 35 °C shows a prominent peak (medium angle peak, M) at 14.369 ± 0.002 nm (mean ± SEM, n= 4), with two smaller satellites at 14.578 ± 0.013 nm (low angle peak, L) and 14.169 ± 0.007 nm (high angle peak, H), for an overall M3 spacing (*S*_M3_) of 14.354 ± 0.011 nm (see also ^3^). *S*_M3_ remains almost constant up to 27 °C and then decreases sharply attaining a minimum of 14.326 nm at 10-15 °C. This reduction (0.2%, Fig. 2g) is similar to that observed at the onset of stimulation, before force develops, both in skeletal^49, 50^ and in cardiac muscle^13^. The M6, originating mainly from a periodic mass distribution in the thick filament backbone with periodicity 7.17 nm^51, 52^, reduces its intensity (*I*_M6_) with cooling (Extended Data Fig. 1f) throughout the whole range of temperatures like M3, but, unlike M3, at temperatures below 27 °C, it is accompanied by the increase in spacing (*S*_M6_, Fig. 2h), which becomes ∼0.25 larger at 10 °C. This change signals the extension of the thick filament, which, in the absence of force, suggests a structural nature as that accompanying thick filament activation^3, 11, 12, 53, 54^. Lowering temperature induces a small but detectable change in the fine structure of the M3 reflection, with an increased separation between the position of the interference peaks indicating a reduction of the interference distance *ID*, the distance between the centres of mass (COM) of the two motor arrays on the thick filament. *ID* can be estimated from the reciprocal of the difference in position between two consecutive peaks, calculated as the reciprocal of the separation between either the M and H peak (*ID*_H_), or the M and L peak (*ID*_L_) (see Supplementary Note 1). In diastole the H peak is the higher satellite peak and *ID*_H_ (Fig. 2i) is preferred as an estimate of *ID*. *ID*_H_ is ∼1020 nm at 35 °C and reduces monotonically attaining ∼760 nm at 10 °C (Fig. 2i). Note that *ID*_H_ is larger by ∼20% than the true distance between the COM of the two motor arrays (see Supplementary Note 1 and Extended Data Fig. 2a-b for the difference between *ID* and that estimated by peak separation). Thus, *ID* is 810 nm at 35 °C, consistent with the contribution of the whole array of 49 layers to *I*_M3_ (red arrow in Fig 2l) and reduces to 760 nm at 27 °C (orange arrow).

In the muscle at rest the myosin motors that leave their ordered resting (OFF) conformation, in which they lie on the surface of the thick filament folded on their tails, enter a conformationally disordered state that gives a negligible contribution to the M3 reflection^40, 48, 55, 56^. Under this condition only the OFF motors contribute to the M3 reflection and *I*_M3_ is approximately proportional to the square of the number of OFF motors^57^. These motors lie along the filament on helical tracks with 43 nm periodicity giving rise to the ML1 layer line, and *I*_ML1_ too is expected to vary with the square of the number of OFF motors. The dependence of the fraction of OFF motors (*f*_OFF_) on the temperature obtained as the square root of either *I*_M3_ (filled circles) or *I*_ML1_ (open circles), normalised for the value at 35 °C, is shown in Fig. 2j. Cooling the muscle down to 10 °C, which reduces both *I*_M3_ (Fig. 2f) and *I*_ML1_ (Fig. 2d) to ∼0.2 the value at physiological temperature, reduces *f*_OFF_ to 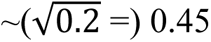.

Given that only the OFF motors contribute to *I*_M3_, the decrease in *ID*_H_ (Fig. 2i) accompanying the reduction of the number of OFF motors can be explained solely by axial broadening of the intensity profile generated by each motor array associated with a shorter coherence length^55, 58^, which implies that the switch of motors from the ordered to the disordered state concerns preferentially motors of the crowns at the two edges of the array (proximal and distal to the centre of the sarcomere)^40, 48, 55^. This suggests that the MyBP-C in the central 1/3 of the half-thick filament has a stabilizing effect on the IHM against the disordering effect of cooling. The idea was tested with a structural model in which the half-thick filament is divided in three structurally distinct regions (Fig. 2l and Extended Data Fig. 3a), proximal (P-zone), layers 1-6, central (C-zone), layers 7-30 with the MyBP-C, and distal (D-zone), layers 31-49^59–61^. The model (for details see Supplementary Note 2) assumes that cooling perturbs, in a different but homogeneous way, the three regions, inducing the release of a different fraction of motors from their OFF state in each of them (Extended Data Fig. 3b). Fitting the intensity profiles of the M3 reflection with this model (Extended Data Fig. 3c-d) predicts that the fraction of OFF motors in the P- and D-zones (Fig. 2k, *f*_OFF,P_, green diamonds and *f*_OFF,D_, brown diamonds, respectively) decreases more than that in the C-zone, *f*_OFF,C_ (black diamonds). In particular, cooling from 35 to 10 °C decreases *f*_OFF,P_ and *f*_OFF,D,_ by 64 ± 7% and 77 ± 4%, respectively, while decreases *f*_OFF,C_ by only 32 ± 8%, so that the overall decrease is by 57% (gray diamonds, predicted *f*_OFF_ = (6·*f*_OFF,P_+24·*f*_OFF,C_+19·*f*_OFF,D_)/49), in agreement with that derived from *I*_M3_ and *I*_ML1_ changes (Fig. 2j, gray dashed line from that in k). At 27 °C, *f*_OFF,P_ and *f*_OFF,D_ are already reduced by almost the same 30% (for the statistics see Supplementary Note 2 and Extended Data Fig. 3b), while *f*_OFF,C_ is still fully preserved (Fig. 2k). A graphical representation of the probability of motors in crowns P, C and D to be OFF and ON at 27 °C is given in Fig 2m by the relative lengths of cyan (*f*_OFF_) and gray (*f*_ON_) bars in each layer.

### Motor attachment upon stimulation starts in the C-zone and spreads out with systolic force

The higher sensitivity of the peripheral regions of the half thick filament to the disordering action of cooling, related to a stabilising action of the MyBP-C on the OFF state of myosin motors^60, 62^, suggests a possible regional hierarchy privileging the distal regions of the thick filament for initial motor attachment to actin and force generation. A crucial temperature to test the idea under both structural and functional points of view is 27 °C, as it is high enough to fully preserve the OFF state in the C-zone, while inducing a ∼30% drop of *f*_OFF_ in the P and D zones (Fig. 2k). Moreover, 27 °C is the near-physiological temperature selected for most of mechanical and X-ray diffraction experiments in intact cardiac preparations^3, 13, 40, 63^.

How the regional differences in the regulatory state of the thick filament at rest contribute to systolic force and its modulation was determined by collecting the relevant X-ray diffraction signals during twitches with different forces at 27 °C. The force at the peak of the twitch (*T*_p_) was varied in the range 5-100 kPa under different protocols that are known to vary the force by varying the number of attached motors (Methods and Fig. 3a-d). For comparison the intensity profiles of the reflections marking the regulatory state of the filament at rest (Fig. 3e-h and Extended Data Fig. 4a, blue) are superimposed on those recorded at the peak of a low force twitch (*T*_p_ 23 kPa, green) and of a relatively high force twitch (*T*_p_ 64 kPa, magenta). The relevant structural parameters are plotted versus *T*_p_ in Fig. 3i-p and Extended Data Fig. 4b. The points are average values obtained by grouping pooled data in five classes (Extended Data Fig. 5, where the limits of each class are indicated by the vertical dashed lines). The point lying on the ordinate axis in panels i, j and l-p of Fig. 3 shows the value at rest obtained by averaging the points on the ordinate in Extended Data Fig. 5, where they are distinct for quiescence (blue) and diastole (cyan) to show that they are not significantly different^3, 40^.

**Figure 3.**
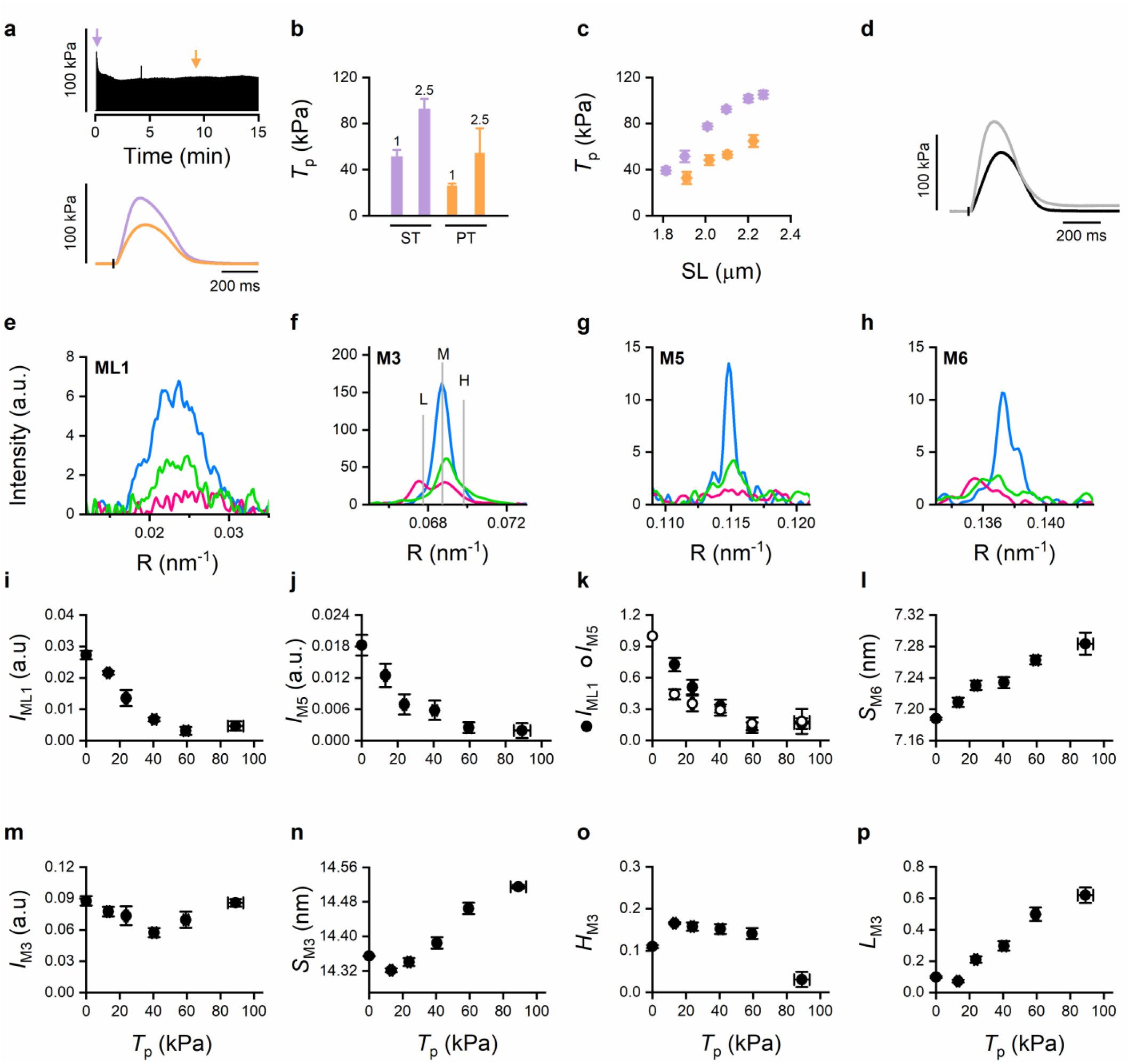
Protocols used to change *T*_p_ and *T*_p_-dependence of the relevant structural parameters marking the regulatory state of the thick filament. **a.** Time course of the *T*_p_ changes following the start of pacing at 0.5 Hz, SL ∼2.1 μm. The arrows indicate the twitches shown superimposed in the lower panel (violet ST, orange PT). **b.** *T*_p_ at different [Ca^2+^]_o_ as indicated by the numbers (mM) above the bars, SL ∼2.1 μm, Violet ST, orange PT. **c.** Force-sarcomere length relation at the peak of the twitch in 1 mM [Ca^2+^]_o_ (violet, ST; orange, PT). Averaged data pooled from twitches in either FE or LC conditions, range of SL before stimulus 1.9 – 2.3 µm. n range 6-12 (PT), 6-51 (ST). **d.** Superimposed force traces for single twitches in FE (black, 2.0 μm SL at *T*_p_, and LC (gray, 2.15 μm SL at *T*_p_). Before stimulus, SL 2.19 μm; [Ca^2+^]_o_ = 1 mM. **e-h**. Superimposed intensity distribution in the reciprocal space (R) of the ML1 (e), M3 (f), M5 (g) and M6 (h) at rest (blue) and at *T*_p_ 23 kPa (green) and 64 kPa (magenta). **i-p.** Plots of the relevant structural parameters versus *T*_p_, with the resting value reported on the ordinate axis: i. *I*_ML1_. j. *I*_M5_. **k.** Superimposed *I*_ML1_ (filled circles) and *I*_M5_ (open circles) changes relative to their resting value. At 13 kPa *I*_ML1_ is 0.73 ± 0.06 and *I*_M5_ is 0.44± 0.05. *P* for the significance of the difference = 0.001 (*df* =4). l. *S*_M6_. m. *I*_M3_ (corrected for the change in the radial width with respect to rest). n. *S*_M3_. o *H*_M3_. p. *L*_M3_. Data are mean ± SEM from pooled data grouped in five classes as detailed in Extended Data Fig. 5: n= 3-28, from 13 preparations. Data in k obtained by averaging points in Extended Data Fig. 5, each point made relative to its resting value. CSA 130,000 ± 88,000 µm^2^ (mean ± SD).

The structural parameters can be grouped in different categories according to their dependence on *T*_p_. The increase in *I*_1,1_/*I*_1,0_ (Extended Data Fig. 4b), indicating motors moving away from the thick filament towards the thin filament, and the increase in *S*_M6_ (Fig. 3l), indicating structural extension of thick filament, much larger than that expected from filament elasticity^11, 54^, progress monotonically throughout the whole *T*_p_ range showing some degree of saturation only at the highest *T*_p_ value. Instead, the changes of *I*_ML1_ (Fig. 3i), marking the helical disposition of motors on the surface of the filament, and *I*_M5_ (Fig. 3j), the intensity of the fifth order forbidden reflection, generated by the perturbation of the three axial layers within the fundamental 43 nm periodicity^46^, drop to a minimum constant value ∼15% the resting value, attained at a force ≤60 kPa, that is ∼1/2 the maximum *T*_p_ value (*T*_p,max_, 110 kPa, ^3, 64^). *I*_M3_ (Fig. 3m) (which at rest originates from the OFF myosin motors and at *T*_p_ is contributed also by motors attached to actin, with their lever arms oriented nearly perpendicular to filament axis^3^) shows a biphasic behaviour: it decreases to a minimum value attained at ∼40 kPa and then recovers up to a value similar to that at rest. Similarly, *S*_M3_ (Fig. 3n), 14.35 nm at rest, shows an early reduction at the very low *T*_p_ (13 kPa) (−0.23%, See Extended Data Table 1) and then a monotonic rise to a value 1% larger than at rest, the same % increase found for *S*_M6_. A reduction in *S*_M3_ similar to that observed here has been found in skeletal and cardiac muscles early after the start of stimulation^13, 50^ and in skinned fibres from skeletal muscle at low concentration of activating Ca^2+^ ^65^. Notably a comparable reduction in *S*_M3_ occurs by lowering temperature (Fig. 2g and ^40, 56^). The fine structure of the M3 changes progressively from that at rest, characterised by a main (middle angle, M) peak and two small satellite peaks on each side (L and H respectively), to that typical of the high force systole^3^, characterised by two main peaks, corresponding to the M and L peaks (Fig. 4a and b). The progression of these changes is defined by the *T*_p_-dependent changes in the fractional contribution to *I*_M3_ of the L peak (*L*_M3_, Fig. 3p). The changes indicate that the population of myosin motors switches progressively from a folded OFF state to a state either disordered or attached to actin with the lever arm tilted nearly perpendicular to the filament axis, with a corresponding shift by ∼10 nm of the centre of its axial mass density projection^66^. Notably at 13 kPa, *H*_M3_ (Fig. 3o), the fractional contribution of the high angle peak to *I*_M3_, increases by 50% (Extended Data Table 1) with respect to the resting value, while *L*_M3_ (Fig. 3p), the fractional contribution of the low angle peak, slightly reduces (−8%, Extended Data Table 1). For the same *T*_p_, *S*_M3_ reduces by 0.23%, while the extension of the thick filament, marked by *S*_M6_ (Fig. 3l) increases by 29% (Extended Data Table 1). This indicates that myosin motors, for the most part still in their OFF state at this low force, have undergone some sort of disengagement from the surface of the thick filament that causes their axial periodicity to reduce more than the overall filament length causing a change in the phase of the interference function that samples the M3 reflection with consequent increase in *H*_M3_ and reduction in *L*_M3_.

**Figure 4.**
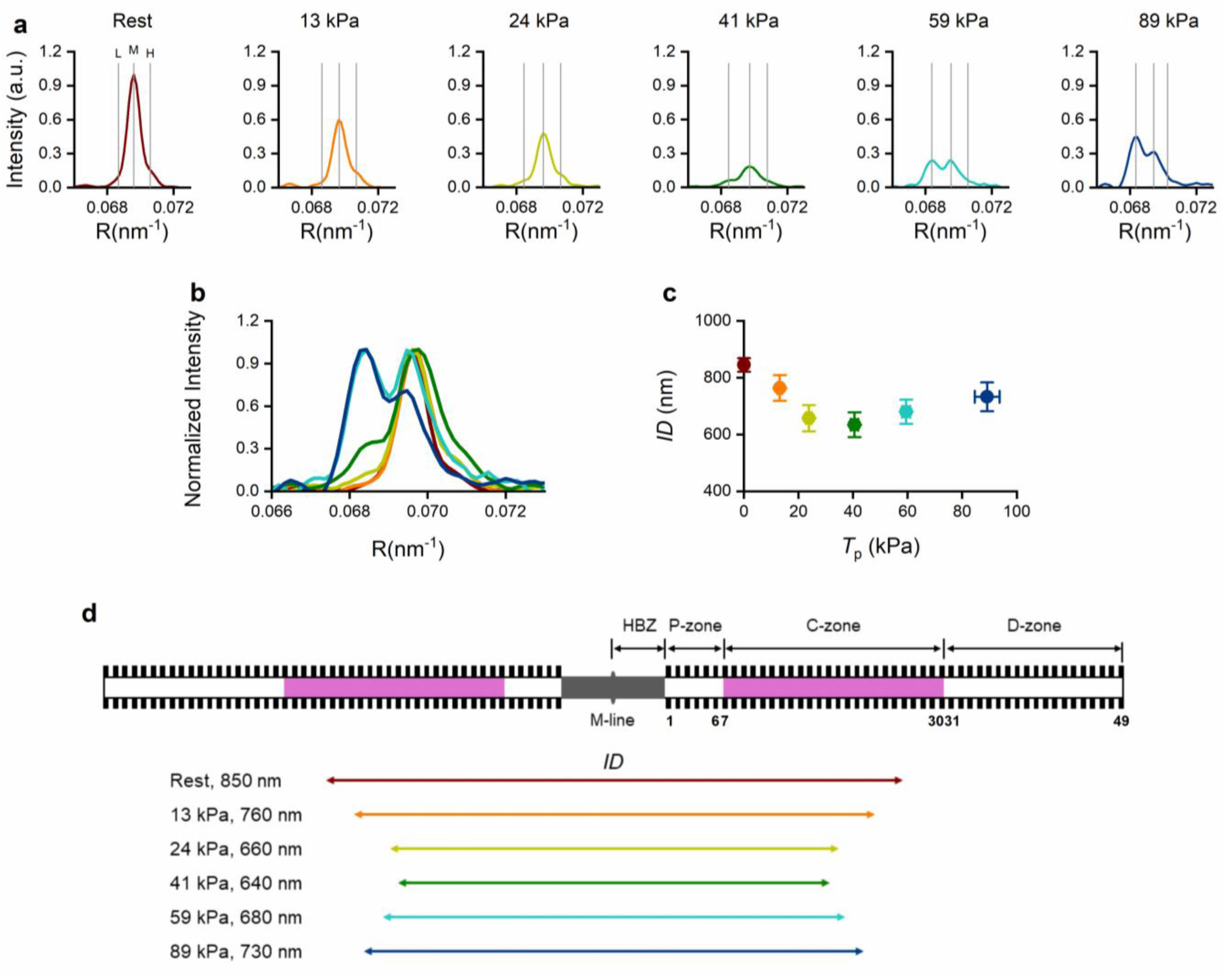
The source of the interference fine structure of the M3 reflection at different *T*_p_. **a.** Axial intensity distribution in the reciprocal space (R) of the M3 reflection at rest and at different *T*_p_ as indicated by the colour code (the same colour code is used also in panels b-d). Vertical lines mark the position of the three component peaks. **b.** Superimposed intensity distributions from a, after normalisation for the intensity of the highest peak (M for rest and *T*_p_ ≤41 kPa, L for *T*_p_ >41 kPa. **c.** *ID* versus *T*_p._ **d**. Pictorial representation (coloured arrows, same colour code as in a) of the *ID* of the two arrays of diffractors estimated from the M3 fine structure at rest and at different *T*_p_ in relation to the location of the two arrays on the thick filament.

In the investigated range of *T*_p_, the changes in the interference distance *ID*, estimated from the peak separation L-M (*ID*_L_) (see Supplementary Note 1 and Extended Data Fig. 2), show a biphasic behaviour. In the first phase of the *ID* – *T*_p_ relation, associated to rise in *T*_p_ up to ∼40 kPa, *ID* decreases from ∼850 nm to a minimum of ∼640 nm, and in the second phase, associated to further rise in *T*_p_, *ID* recovers up to ∼730 nm (Fig. 4). The biphasic dependence of *ID* on *T*_p_ is parallel to the biphasic *T*_p_- dependence of *I*_M3_. For *T*_p_ <40 kPa, both *I*_M3_ and *ID* reduce, suggesting that in this range of forces the M3 reflection is dominated by the decreasing contribution of OFF motors in the distal region of the thick filament. In contrast, for *T*_p_ >40 kPa, both *I*_M3_ and *ID* increase, suggesting that in this range of forces the dominant contribution to the M3 is given by the fraction of attached motors increasing progressively from the centre to the periphery of the thick filament. The idea that motors switch from the OFF state to the disordered state within a relatively narrow range of forces (<60 kPa) is further supported by the finding that *I*_ML1_, originating solely from the helical symmetry of motors in the OFF state, saturates at a minimum value at *T*_p_ ≤60 kPa.

The *ID* changes with *T*_p_ are represented in Fig 4d by the length of the arrows reporting the centre-to-centre distance of the two arrays of motors on the thick filament, assuming the start of the array to coincide with the first layer. The proper definition of the zonal distribution of layers contributing to *I*_M3_ at each *T*_p_ requires the best fitting of the whole set of parameters from Fig. 3 with a structural model simulation (see next section).

For *T*_p_ > 40kPa, at which the drop in *I*_ML1_ (Fig. 3i) and thus the process of motor switching OFF are almost complete, *ID* increases in proportion to *T*_p_ (Fig. 4c) and number of attached motors. This is a clear indication that further attachments concern progressively more peripheral regions of the thick filament. These regions, however, are activated also at lower values of *T*_p_. In fact, in the *T*_p_ range <40 kPa, the simultaneous *T*_p_-dependent drops of both *I*_M3_ (Fig. 3m) and *ID* (Fig. 4c) clearly indicate that the decrease of population of ordered folded OFF motors occurs mainly at the periphery of the filament. Thus, at odds with the original mechanosensing concept^3^, for *T*_p_ >40 kPa the increase of attachments with the increase in systolic performance is not accounted for by switching ON of motors. In the cardiac systole the intracellular [Ca^2+^] rises transiently to a level (∼10^-6^ M) lower than that for full thin filament activation. At this Ca^2+^ concentration the Tm has only partially undergone the azimuthal movement that underpins the progression of the thin filament through the transition that makes the actin sites available for interaction with myosin^8, 30^. Namely, of the three states identified by the steric-blocking model (blocked, closed and open), the thin filament occupies mainly the blocked and closed states, a condition that makes the filament quite sensitive to cooperative activation by myosin motors^38, 44, 67^. In the presence of a fully active thick filament in the P- and D-zones and under the condition that low forces are accounted for by motors attached to actin from the crowns in the C-zone, the cooperative action of these motors on thin filament activation can *per se* explain why the progression of attachments with force occurs toward the periphery of the half-thick filament. The attached motors from the crowns at the end of the C-zone make actin monomers available for attachment of motors from the crowns in the neighbouring D- and P-zones. These new attachments in turn activate neighbouring actin monomers, allowing attachments from progressively more peripheral crowns.

### Structural model simulation of thick filament activation and motor attachment

A structural model simulation of the changes of the M3 intensity distribution with *T*_p_ is used to define a regional hierarchy of motor activation along the thick filament and select between three alternative hypotheses on the localization of attached motors and their progression with force. The first hypothesis (Model 1) is that motors attach first in the C-zone and further attachments with increase in *T*_p_ progress towards the periphery of the half-thick filament. The second hypothesis (Model 2) is that motors attach uniformly throughout the 49 crowns of the half-thick filament and increase in *T*_p_ is achieved with homogeneous increase in the fraction of attached motors in each crown. The third hypothesis (Model 3), suggested by the original concept of thick filament mechanosensing, assumes that first motor attachments occur at the edges of the thick filament and then progress towards the centre of the thick filament with the increase in *T*_p_.

The starting point of the model simulation sets the fraction of motors in the OFF conformation in P-, C- and D-zones as predicted at rest at 27 °C (Fig. 5a, rest): *f*_OFF,C_ = 1, *f*_OFF,P_ ≡ *f*_OFF,D_ = 0.71 (see Supplementary Note 2). With respect to the simulation used to establish the effect of temperature on the OFF state of the motors, the simulation at *T*_p_ is complicated by the requirement to consider the contribution of the attached motors and their detached partner in the dimer^68, 69^, the COM of which differs from that of the dimer in the OFF state by 10.5 nm away from the centre of the filament^3, 66^. We indicate with *f*_A_ the fraction of attached motors in each crown of 6 motors contributing to M3. With *n* contributing crowns in the half thick filament, the total number *N*_A_ of attached motors in each half thick filament is *n*·6·*f*_A_. Given the protocols used to vary *T*_p_ (see Methods and ^63^), a general constraint of model simulation is that *N*_A_ scales linearly with *T*_p_. At the maximal systolic force *T*_p,max_ (110 kPa, ^3^), at which *f*_A_= 0.22 and *n*= 49, *N*_A_ is 65 ^3^. At any force the attached motors are contributed from layers between *n*_i_ and *n*_e_, the proximal and distal layers relative to the M-line, with *n*_i_=1 and and *n*_e_= 49 at *T*_p,max_. Thus *f*_A_·(*n*_e_ – *n*_i_ +1), that is *N*_A_/6, scales linearly with *T*_p_.

**Figure 5.**
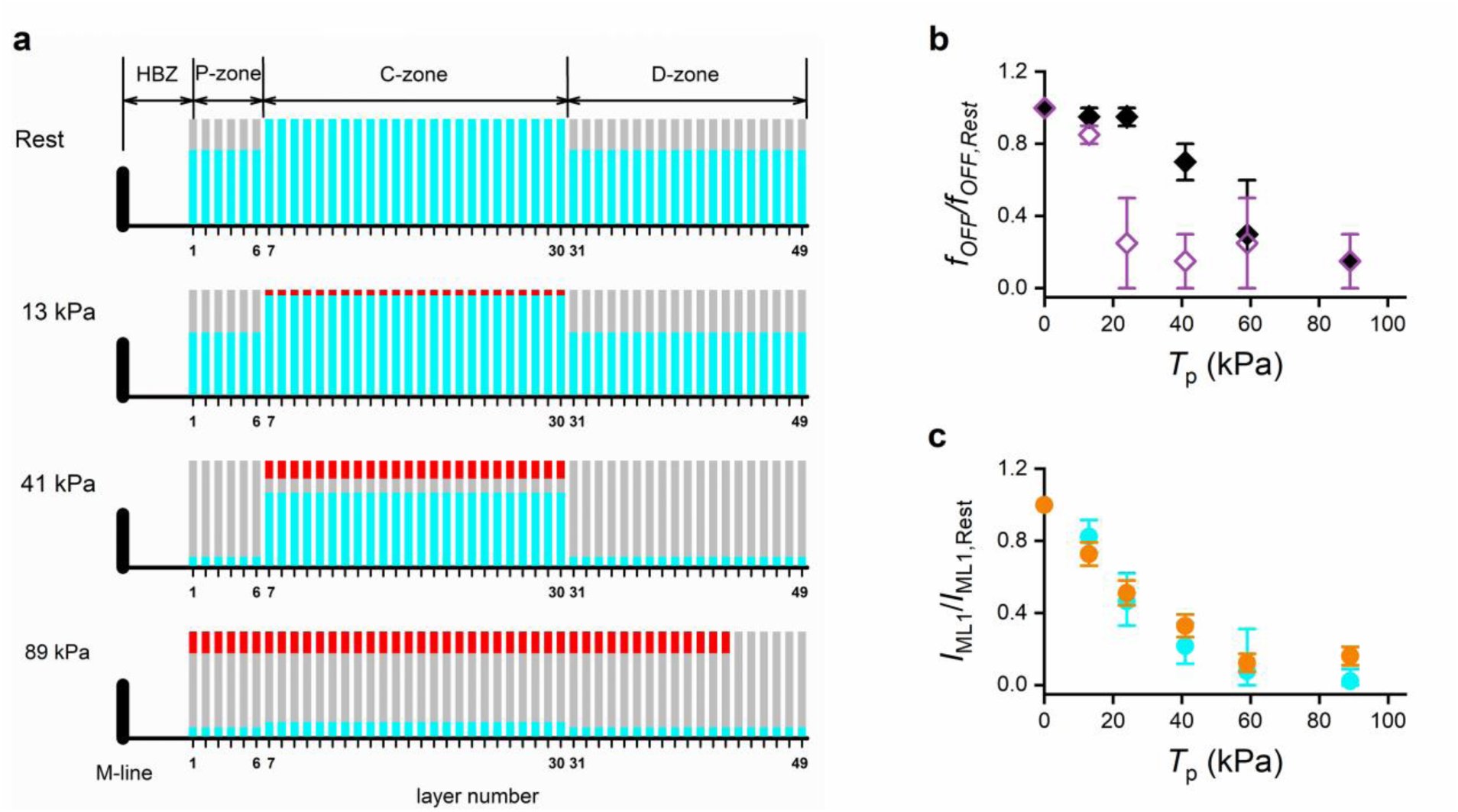
Model 1 simulation of motor fractional occupancy of the OFF, ON, and attached state. **a.** Schematic representation of the probability of each layer along the half-thick filament to have motors in the OFF state (*f*_OFF_, cyan) or in the ON state (*f*_ON_, gray) at 27 °C at rest (1^st^ panel) and at *T*_p_ 13 kPa (2^nd^ panel), 41 kPa (3^rd^ panel) and 89 kPa (4^th^ panel). Motor layers are numbered 1-49 starting from the centre of the thick filament (M-line). P-zone, layers 1-6; C-zone, layers 7-30; D-zone, layers 31-49. The fraction attached motors (red) is defined according to the assumptions of model 1 (see text). The three fractional occupancies in each layer are represented through the length of the corresponding color in the bar. **b.** Model 1 prediction of the *T*_p_-dependence of the fraction of motors in the OFF state relative to the fraction at rest at 27 °C within the C-zone (filled diamonds) and outside the C-zone (open diamonds). **c.** Superimposed I_ML1_ – *T*_p_ relations: orange, observed (from Fig 3i); cyan, predicted by Model 1.

The best structural model is selected on the basis of its ability to simulate the relevant parameters that characterise the M3 intensity distribution: *I*_M3_, *S*_M3_, *L*_M3_ and *ID*_L_. For any given *T*_p_ (first column in Supplementary Tables 1-3) and number of attached motors (second column) the model output is calculated, after adjustment of the axial periodicity of diffractors (*d*_0_) to get the best fit of *S*_M3_, for pairs of progressively reduced *f*_OFF,C_, *f*_OFF,D_ values (with *f*_OFF,P_ the same as *f*_OFF,D_). Details of the procedure are given in Supplementary Note 3. An [*f*_OFF,C_; *f*_OFF,D_] pair is considered adequate to reproduce the data when the model output with that pair gives estimates of the three parameters (*I*_M3_, *L*_M3_, *ID*_L_) falling inside the range of ±2·SEM (p<0.05) from the observed mean values. Only Model 1 (Fig. 5, Extended Data Fig. 6 and 7 and Supplementary Table 1) holds values for *f*_OFF,C_ and *f*_OFF,D_ that allow simultaneous reproduction of the observed *I*_M3_, *L*_M3_ and *ID*_L_ with the required approximation for all *T*_p_ values. Model 2 (Extended Data Fig. 8 and Supplementary Table 2) fails to reproduce the observed data with the required approximation for forces >24 kPa and Model 3 (Extended Data Fig. 9 and Supplementary Table 3) fails to predict the observed data for all *T*_p_ values. The possibility of Model 2 to reproduce the M3 fine structure at very low forces is related to the reduction of the weight that so few attached motors have on shaping the M3 reflection. However, also with the limited contribution of attached motors at low forces, Model 3 fails at all *T*_p_, demonstrating the inconsistency of the assumption that first motor attachments occur from the edges of the thick filament.

A most important prediction of Model 1 is the zonal difference of the effect of stress on the OFF-ON motor switch. Switching ON of motors in the P- and D-zones is completed at *T*_p_ < 40 kPa (Fig. 5b, open symbols)), while in the C-zone it is completed at *T*_p_ ≥ 60 kPa (filled symbols).

Even if the structural model is used to fit the M3 reflection, its prediction for *f*_OFF,C_ and *f*_OFF,D_ (=*f*_OFF,P_) gives an overall fraction *f*_OFF_ of OFF motors that satisfactorily accounts for the drop in *I*_ML1_, as demonstrated by superimposing the observed *I*_ML1_-*T*_p_ relation (Fig. 5c, orange, from Fig. 3i) to the relation obtained by squaring *f*_OFF_ (cyan).

## DISCUSSION

### Matching X-ray diffraction signals with the structure of the thick filament

The cryo-EM 3-D description of the architecture of cardiac thick filament^4, 5^, showing the details of the interactions between titin, MyBP-C and myosin heads and tails, provides the framework for a molecular interpretation of the present results. The lead over filament stress of the process of motor switching ON throughout the whole thick filament, as shown by the relation between *I*_ML1_ and *T*_p_, calls for a role of A-band titin in thick filament activation. In fact, as suggested by the EM structure of the P- and C-zone^4, 5^, titin is able to mechanically control the thick filament structure all over its length via its extended interactions with myosin tails within each 43 nm triplet of crowns working as individual cooperative units. The idea of titin as the molecular player for thick filament mechanosensing that switches motors ON for forces < 60 kPa is solidified by the finding that I-band titin effective stiffness increases by two orders of magnitude upon stimulation of the skeletal muscle^6^, enabling titin to efficiently transmit any mechanical/structural perturbation to the thick filament and specifically affect the helically ordered motor disposition^6^. The *T*_p_ dependence of the intensity of the forbidden reflections originated from the triplet perturbation with 43 nm fundamental periodicity (in Fig. 3g the fifth order, M5) further strengthens this idea. In fact, M5 as well as the high angle component of the M2 share an interference fine structure (see Fig. 2b and c) consistent with their origin from a perturbing structure with fundamental 43 nm axial periodicity extending throughout most of the thick filament (Extended Data Fig. 10 and 39^,40^), that is likely titin. Consequently, the finding that at the smallest *T*_p_ (13 kPa), with respect to the value at rest, *I*_ML1_ decreases by 27%, while *I*_M5_ decreases by 56% (Fig. 3k) does indeed support the idea that an activation-dependent titin change (question mark in Fig. 6) triggers the loss of the helically ordered conformation of myosin motors. This conclusion is strengthened by the finding that in stimulated fibres from skeletal muscle with myosin motors activation pharmacologically suppressed, a force step causes the drop in *I*_M5_ and *I*_ML1_ with a similar temporal hierarchy (see Fig. 4 in ^6^): while *I*_M5_ drops simultaneously with the imposed filament strain (estimated by the increase in *S*_M6_), *I*_ML1_ drops with a slower, exponential time course. Other X-ray diffraction signals supporting the role of A-band titin as a trigger for motor switching ON following stimulation are the peculiar changes in spacing and fine structure of the M3 at the smallest *T*_p_ (13 kPa, see Extended Data Table 1). The reduction of *S*_M3_ (Fig. 3n) occurs while the thick filament extension, marked by *S*_M6_ (Fig. 3l), increases, and is accompanied by the increase of *H*_M3_ (Fig. 3o) and reduction of *L*_M3_ (Fig. 3p). Assuming that at 13 kPa, for the paucity of attached motors (8 per half thick filament, Supplementary Table 1), all these signals are dominated by the contribution of motors in the OFF state, their changes indicate a disengagement of the motors from the ordered folded disposition on the surface of the filament caused by the disruption of the triplet axial perturbation by activated titin (Fig. 6).

**Figure 6.**
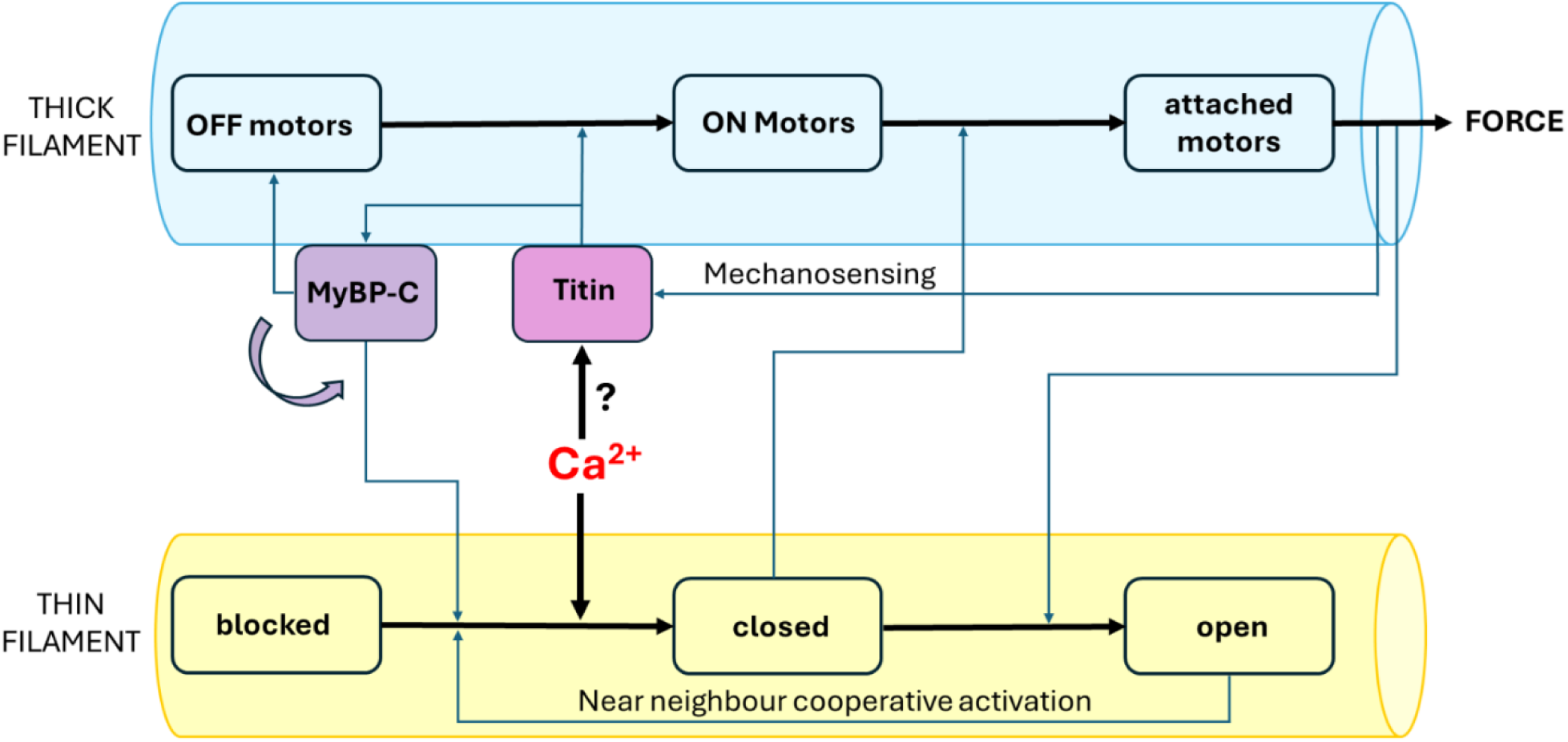
Interactions among titin, MyBP-C and myofilaments and their function in the regulation of the cardiac output. Rise in internal Ca^2+^ triggers activation of the thin filament (yellow) and, through a still undetermined process, activation of titin (magenta) that switches ON the thick filament (cyan) and then the N-terminus of MyBP-C (violet) accounting for initial availability of actin for motor attachments from the C-zone. The ensuing force development and titin-mediated mechanosensing complete the thick filament activation. At subsaturating level of internal Ca^2+^ near neighbour cooperative activation of thin filament by attached motors allows further attachments from the more peripherical regions of the half thick filament.

The switching ON of motors is hierarchically distributed along the thick filament, as demonstrated by the early reduction of *ID* at *T*_p_ <40 kPa, reproduced by Model 1 (Fig. 5a, second and third panel, and Extended Data Fig. 5b-d) with a selective switching ON of P- and D-zones, shown by gray (ON) growing at the expenses of cyan (OFF). This characteristic of thick filament activation contrasts with the original concept of thick filament mechanosensing and is in accordance with the larger propensity of motors to be switched ON in the D- and P-zones not only by increase in stress but also by lowering temperature (Fig. 2m). The higher resistance of the OFF motors in the C-zone against disordering factors points toward a stabilizing effect of MyBP-C on the IHM. This conclusion is consistent with the cryo-EM demonstration of the specific interactions of the C-terminus of MyBP-C with the myosin IHM and tails^4, 5^: in more detail, the C-terminus of MyBP-C binds from one side the tails of three consecutive crowns of type D (in the nomenclature of ^4^), thus spanning its effect throughout three 43 nm periods, and from the other side the IHM of crowns T and H within the same 43 nm period. Moreover, the N-terminus of MyBP-C, when bound to the head-tail junction of the myosin motors, could contribute to IHM stabilisation^18, 42, 70^.

At odds with the stabilising effect of MyBP-C on the OFF structure of thick filament, Model 1 indicates that upon stimulation the crowns contributing to motor attachments at *T*_p_ ≤40 kPa are limited to the C-zone of the thick filament (Fig. 5a, second and third panel, and Extended Data Fig. 6b-d) with attached motors (red) growing at the expenses of OFF motors (cyan). Thus, the most resistant zone to switching motors ON by the perturbing action of cooling and stress contributes the first motor attachments. In contrast to the classical concept of mechanosensing, this result indicates that switching motors ON is not *per se* sufficient to promote attachments, while MyBP-C should serve a specific role in triggering the first attachments, through its C-terminus binding the myosin heads and tails and its N-terminus switching from binding the myosin head-tail junction to binding actin (Fig. 6, violet arrow) Both EM and fluorescent labels demonstrated that at low [Ca^2+^] the MyBP-C N-terminus binds actin near the interaction site of Tm^7, 41–43^, displacing Tm from the position that is inhibitory for motor attachment (blocked, ^8, 30^) towards the position that allows motor attachment. This activating effect of the N-terminus is confirmed by the increase of Ca^2+^ sensitivity in filament motility assay^7, 43^.

In the cardiac systole internal [Ca^2+^] rises to ∼10^-6^ M, a concentration low enough to preserve a relatively high probability for the regulated actin filament to be in the blocked state, so that thin filament activation is quite sensitive to cooperative action of myosin motors^38, 44, 67^. Under this condition, the efficiently fast force development at the start of contraction, when there are no attached motors, calls for the presence of a supplementary mechanism of thin filament activation. The finding in this work that low *T*_p_ twitches (<40kPa) are accounted for by actin-attachment of motors from crowns in the C-zone, strongly supports the idea that the N-terminus of MyBP-C surrogates the cooperative action of myosin motors in triggering thin filament activation by switching interaction from myosin head-tail junction on the thick filament to actin on the thin filament^18, 42^. The switch is likely caused by changes in the thick filament structure initiated by Ca^2+^ dependent activation of titin, through a still unknown mechanism (question mark in Fig. 6). The role of titin as a trigger of thick filament activation and ensuing switch of MyBP-C N-terminus is supported by the early drop in titin-dependent triplet axial perturbation of crowns. Notably, the action of MyBP-C N-terminus as thin filament activator is restricted to relatively low Ca^2+^ concentrations^7, 41, 43^, which makes this mechanism peculiar for the heart, in which the rise in internal [Ca^2+^] during the systole is limited to ∼10^-6^ M. This mechanism may be even more capital in cardiac than in skeletal muscle, given the lower affinity of cardiac TnC for Ca^2+^ ^7^.

The relatively low internal [Ca^2+^] in the cardiac systole keeps high the probability for a thin filament regulatory unit to be in the blocked state, making the cooperative activation of thin filament by motor attachment the driving mechanism for further attachments^8, 44^. For forces >40 kPa, even if the switching ON of motors in P- and D-zones is complete (Fig. 5a, 41 and 89 kPa and Extended Data Fig. 5d-f, where gray almost fully replaces cyan, and Supplementary Table 1), the increase in attachments (red) with the increase in *T*_p_ is characterised by a progressive spread toward the periphery. This prediction of Model 1 is fully accounted for by near-neighbour cooperative thin filament activation (Fig. 6): motors attached from the crowns at the end of the C-zone make actin monomers available for new attachments from the crowns in the neighbouring D- and P-zones. These new attachments in turn activate further peripheral actin monomers, allowing attachments from progressively more peripheral crowns of the thick filament. The prediction from Model 1 that, at 89 kPa, attached motors span only up to crown 43 is explained by considering that *T*_p_ is 20% lower than the maximum systolic force (110 kPa, ^3^) attained with attached motors spanning all the 49 crowns.

### Physiological implications

During the systole the heart pumps the blood into the arterial system with a power that adapts to the load, namely the arterial pressure, in order to maintain the flux through the circulatory system constant. According to the classical Frank-Starling law the mechanism of adaptation implies that the strength of the contraction in the systole increases with the amount of blood filling the ventricles during the preceding diastole. An increase in the arterial pressure that reduces the stroke volume causes an increase in the volume attained at the end of the subsequent diastolic refilling, which induces a stronger contraction in the next systole. The equivalent of the Frank-Starling law at the level of the sarcomere is the LDA^35, 36^, according to which the relation between SL and active force is much steeper than that of the skeletal muscle for the same degree of filament overlap. The physiological level of internal Ca^2+^ concentration attained during the systole is limited to 10^-6^ M, thus other fundamental players in cardiac contractility are Ca^2+^ sensitivity of the thin filament and cooperative activation of the filament by attached myosin motors. Both LDA and Ca^2+^ sensitivity in demembranated cardiac myocytes were found to be enhanced by the degree of phosphorylation of accessory sarcomeric proteins like MyBP-C^18, 20, 21, 32, 33^. On the whole, the idea emerging from those studies was that titin-mediated stress on the thick filament and the degree of MyBP-C phosphorylation exert their positive inotropic effect by increasing thick filament activation during the diastole.

This idea, however, was contradicted by X-ray diffraction experiments on electrically paced intact trabeculae from the rat ventricle, demonstrating that the degree of activation of the thick filament is tuned to the loading conditions of the systole, which determine the end systolic SL, independently of the diastolic SL^3, 63^ and that the positive inotropic action of protein phosphorylation does not affect the folded, helically ordered state of motors in diastole^37^. In addition, the classical interpretation of the LDA as due to the increase in titin-mediated stress on the thick filament with the increase in SL in diastole was weakened by the experimental evidence that in the range of physiological SL (1.9-2.2 μm) there is no structural signal of motors leaving the OFF state^37^. A further element that undermines a titin-mediated activation of thick filament in diastole is the evidence from skeletal muscle that, within the physiological SL (2-2.6 μm in that case), I-band titin becomes an efficient coupler for the transmission of the stress only upon activation^6^.

How do the results presented here and their interpretation in light of recent Cryo-EM description of thick filament structure^4, 5^ fit into the current views on the mechanism of regulation of systolic performance? The first question to be taken into account is the definition of the mechanism controlling the regulatory state of the thick filament, namely the fraction of switched ON motors throughout the contraction cycle. In the intact myocyte at physiological temperature there is no evidence for a significant fraction of constitutively ON motors present at rest either in skeletal muscle^56^ or in the heart (Fig. 2). Instead, the plethora of extended interactions of the A-band titin with the myosin tails within each 43 nm triplet of crowns^4, 5^, integrated with the activation dependent stiffening of I-band titin^6^, provide new and solid molecular basis for interpreting the early switching ON of myosin motors and its regional characterization along the thick filament in relation to force. At the lowest *T*_p_ (13 kPa), when most of motors are in the OFF state (Fig. 5b, Supplementary Table 1), the reduction of axial motor periodicity (*S*_M3_, Fig. 3n) and the increase in filament extension (*S*_M6_, Fig. 3l), together with the changes in M3 fine structure (increase of *H*_M3_, Fig. 3o, and reduction of *L*_M3_, Fig. 3p) and the loss of triplet axial perturbation (reduction of *I*_M5_, Fig. 3j) indicate that myosin motors have already undergone a disengagement with respect to their resting, folded disposition on the surface of the thick filament. The drop in the intensity of M5, the interference fine structure of which reveals its origin from a perturbing structure covering almost the whole filament like titin (Extended Data Fig. 10), precedes the drop of *I*_ML1_ and thus of the ordered helical disposition of OFF motors (Fig, 3k), specifically indicating that is the Ca^2+^-dependent activation of titin^6^ that underpins the mechanosensing-based process that switches myosin motors ON (Fig. 6). Switching ON progresses from the periphery of thick filament, where it saturates at forces ≤40 kPa (Fig. 5b, open diamonds), to the C-zone where saturation is attained only at forces ∼60 kPa (filled diamonds). Thus, titin-mediated thick filament activation is less effective in the C-zone due to the specific stabilising action of the MyBP-C on the IHM, but in any case attains saturation at forces ∼½ the maximum systolic force (Fig. 5c). This conclusion limits the effect of mechanosensing-based regulation of thick filament on the systolic performance to forces ≤1/2 the maximum force.

The second question to be taken into account is an explanation for the regional hierarchy of motor attachment along the thick filament. At odds with earlier switching ON of motors at the periphery of the thick filament, first motor attachments occur in the C-zone, a result that imposes a revision of the idea that attachment must be preceded by disordering of motors. An explanation is that MyBP-C N-terminus drives in some way myosin motors to attach to actin by switching interaction from myosin head-tail junction on the thick filament to actin on the thin filament^18, 42^, a process triggered by early structural changes in the thick filament C-zone initiated by Ca^2+^ dependent activation of titin (Fig. 6). A final question concerns the need to clarify the relation between the parameters underpinning the systolic performance at the organ level and in a trabecula or a papillary muscle, which are made by columns of myocytes generating stress along their axis. The demonstration that during systole the thick filament is fully switched ON at forces ∼60 kPa (1/2 the maximum force) defines two regimes in relation to the systolic performance. In the range of forces <60 kPa it is the load that defines the energetic cost of the systole since the fraction of motors switching ON depends on the stress on the thick filament. At forces >60 kPa, at which the thick filament is almost fully active, it is the strain dependent kinetics of the motors that define the energetic cost of the performance. In this regime, the load determines the kinetics of the interactions, that is at higher load the rate of motor detachment is slower and the duty ratio and thus the fraction of attached motors is larger.

The heart ventricles work with afterloaded contractions, pumping blood into the vessels once the isovolumetric contraction has attained the arterial pressure. During the systole the pressure in the left ventricle attains ∼120 mmHg, that is ∼16 kPa. The corresponding average load experienced by the myocytes in the ventricle depends on the shape of the ventricle and the thickness of the wall with respect to the cavity volume and for the healthy heart it has been calculated to be not more than three times higher than the pressure (≤ 50 kPa, ^72^). In this respect it must be noted that the wall stress could be an underestimate of the actual force exerted by myocardial fibres, depending on the dispersion of myofibre orientation, including fibres at angles out of the plane of the wall^73^. Keeping the simplest original approach, we can conclude that during the systole the myocytes work just within the force range (60 kPa) for which the thick filament activation is controlled by the load through the titin based mechanosensing. Under this condition the power of the cardiac pump is tuned to the load with maximization of the efficiency. However, under various pathological conditions and in aging, the ratio between wall stress and ventricle pressure may change^74–76^. Expectedly, in Dilated Cardiomyopathy, DCM, thinning of the wall leads myocytes to experience forces > 60 kPa, at which thick filament is fully active, with increase in the energetic cost of the systole. In this case the increase in force is accounted for by further motor attachment from the periphery of thick filament, guided by thin filament cooperative activation.

High spatial resolution X-ray diffraction from the intact multicell preparations (trabeculae and papillary muscles) used in this work is a unique tool to provide molecular signals from the native filament lattice in relation to the mechanical performance. The recent EM description of the packing of the myosin tails and of the interactions between myosin, titin and MyBP-C in the C- and P-zones of the cardiac thick filament^4, 5^ gives an unprecedented tool to understand the X-ray diffraction signals in terms of the role of each molecular player in tuning cardiac performance. Eventually, the advancement in describing the function of each player in the integrated dual filament regulation of the heart output is a prerequisite for its targeting with therapeutic interventions whenever its mutation is causative of a myopathy.

## METHODS

### Sample preparation and stimulation protocols

Male rats (*Rattus norvegicus,* strain Wistar Han, weigh 250–350 g; Charles River, Research Models and Services) were housed at the Bio-Medical Facility (BMF) of the ESRF under controlled temperature (20 °C ± 1 °C), humidity (55 ± 10%), and illumination (12-h light/dark cycles). Food and water were provided ad libitum. Rats were chosen at random for each experiment and sacrificed in agreement with the Italian regulation on animal experimentation (Authorization 17E9C.N.CLU in compliance with Decreto Legislativo 26/2014, following European Community Council Directive 86/609/EC). Rats were anesthetized with isoflurane [5% (vol/vol)], the heart was rapidly excised, placed in a dissection dish and cannulated via aorta and then retrogradely perfused with a modified Krebs–Henseleit (K-H) solution (NaCl, 115 mM; KCl, 4.7 mM; MgSO_4_, 1.2 mM; KH_2_PO_4_, 1.2 mM; NaHCO_3_, 25 mM; CaCl_2_, 0.5 mM; glucose, 10 mM) containing 20 mM 2,3-butanedione monoxime (BDM) and oxygenated with 95% O_2_ and 5% CO_2_ (pH 7.4). Thin, unbranched, and uniform trabeculae or papillary muscles were dissected from the right ventricle under a stereomicroscope (Zeiss SV11 or Stemi 508) by using small knives and scissors at room temperature. Because of its thickness, the preparation was mounted just taut (length *L*_t_) in a dummy trough where it was possible to rotate it along its axis, and the larger and smaller diameters (*D*_l_ and *D*_s_, respectively) were measured using an eyepiece with a graduate scale at three positions along it. The cross-sectional area (CSA) was calculated as *D*_l_ × *D*_s_ × π/4. The dimensions of the preparations (8 trabeculae and 11 papillary muscles) ranged as follows (mean ± SD in brackets): trabeculae, *D*_l_, 320–970 μm (530 ± 220 μm); *D*_s_, 160–320 μm (205 ± 55 μm); CSA, 45,200–137,500 μm2 (83,000 ± 33,000 μm^2^); *L*_t_, 2.3–4.8 mm (3.5 ± 0.8 mm); papillaries, *D*_l_, 300–970 μm (580 ± 210 μm); *D*_s_, 240–620 μm (410 ± 120 μm); CSA, 56,000–426,000 μm^2^ (200,000 ± 120,000 μm2); *L*_t_, 2.0–5.1 mm (3.5 ± 1.1 mm).. The sample was then transferred into a thermoregulated trough (temperature 27 °C unless otherwise specified) perfused (1.2 mL/min) with oxygenated K-H solution and attached via titanium double hooks to the lever arms of a strain-gauge force transducer (AE801, Kronex Technology Corporation, CA, USA) and a loudspeaker motor^77^ mounted on micromanipulators. The preparation was oriented to have the larger transverse axis orthogonal to the X-ray path by twisting opportunely the double-hook arms. A pair of mylar windows bringing two platinum wire electrodes was positioned close to the sample, about 1 mm apart, to minimize the X-ray path in the solution. The trough was sealed to prevent solution leakage and mounted in the path of the X-ray beam at the ID02 beamline of the ESRF^78^, with the long axis of the sample vertical to exploit the smaller size of the beam and allow a higher spatial resolution along the meridional axis of the pattern (parallel to the sample long axis) to better resolve the fine structure of the intensity distribution. The K-H solution perfusing the preparation was exchanged with a K-H solution without BDM and containing CaCl_2_ (either 1 or 2.5 mM). Samples were electrically stimulated via the platinum electrodes with a constant-voltage pulse generator able to deliver pulses of alternate polarity of amplitude up to 30 V. A squared pulse of 1 ms duration was delivered to elicit a single twitch. A stimulus intensity 1.5 times the intensity for the maximal response was selected. At the beginning of each experiment the sarcomere length (SL) was measured by recording sarcomere reflections with a sample-to-detector distance 31 m ^3^ and set at 2.1 μm adjusting the length of the preparation by means of the motor micromanipulator that allowed to measure the change in length of the sample and thus to correct its CSA assuming constant volume behaviour. Two stimulating protocols were used: in the single twitch (ST) protocol single pulses were delivered at the minimum interval (2.5-5 min) necessary to elicit a twitch with a force peak (*T*_p_) that did not further increase by increasing the interval; in the paced twitch (PT) protocol stimuli were delivered continuously at 0.5 Hz frequency (see Fig. 3a). Force, motor lever position and stimulus signals were recorded with a multifunction input/output board (PXIE-6358; National Instruments).

### Mechanical protocols

*T*_p_ was modulated by several protocols that are known to change *T*_p_ by changing the underlying number of attached motors^63^. The protocols concerned either the ST or the PT and included: (*i*) change in the extracellular [Ca^2+^] from 1 to 2.5 mM (Fig. 3b); (*ii*) change of SL in the range 2.0-2.3 µm (Fig. 3c); (*iii*) switch in the mode of operation of the loudspeaker servosystem from fixed-end (FE), in which the trabecula shortens against the compliance of its attachments during force development, to sarcomere length-clamp (LC), in which SL shortening during force development is prevented by feeding the system with a signal based on the changes in SL of a preceding FE twitch.

### X-ray data collection

The X-ray beam at ID02 provided up to 2·10^13^ photons per second at 0.1 nm wavelength, with a size ∼300 μm (horizontal; Full Width at Half Maximum, FWHM) and ∼50 μm (vertical) at the sample. X-ray diffraction patterns were recorded using the FReLoN CCD-based detector with 2,048 × 2,048 pixels (50 × 50 mm^2^ active area and 24 μm pixel size; ^78^). The point spread function (PSF) of the detector was ∼44 μm FWHM. Pixels were binned by eight in the equatorial direction (perpendicular to the sample long axis) before the readout to increase the signal to noise ratio (S/N). The detector was mounted on a wagon inside an evacuated tube 34 m long and 2 m in diameter. The high collimation of the beam allows varying the sample-to-detector distance in the range 0.6-31 m with only a small change of the beam dimensions on the detector making it possible to record both nanometre-scale reflections from the protein periodicities along the myofilaments and micrometre-scale reflections from the sarcomere periodicity^78^. The beam was attenuated to 3% (25-μm thick Fe attenuator) for sample alignment. To minimize radiation damage, X-ray exposure was limited to the data collection period using a fast electromagnetic shutter (model LS200; nmLaser Products, Inc.), and the sample was shifted along its axis by 100 - 200 μm between exposures.

In the experiments on the effects of temperature, SL was preliminarily set to 2.1 μm with sample-to-detector distance 31 m and beam intensity attenuated; then signals up to the M6 reflection were collected with the sample-to-detector distance 1.6 m and full beam intensity. Temperature was changed among 35, 27, 20, 15 and 10 °C in a random sequence, different for the different samples. At each temperature, two patterns were collected with 10-ms exposure from the sample at rest. Collection of patterns at 27 °C was repeated 2-3 times between the other temperatures to monitor for pattern deterioration due to radiation damage.

In the experiments at different *T*_p_, patterns were acquired at rest and at the twitch peak with full beam intensity, with both 31 m (2 ms exposure) and 1.6 m (5-20 ms exposure) sample-to-detector distances. The experiment was stopped when development of contracture or deterioration of the diffraction pattern signalled radiation damage. SL at the peak of the twitch (sample-to-detector distance, 31 m) was measured by using the second order sarcomeric reflection^3^.

### X-Ray data analysis

X-ray diffraction data were analysed using Fit2D (A. Hammersley; ESRF), Sigmaplot (Systat Software Inc.), IgorPro (WaveMetrix Inc.) and Origin-Pro 2021 (OriginLab Corp.).2D patterns were centred and aligned using the equatorial 1,0 reflections and then quadrant folded to increase the S/N. The distribution of diffracted intensity along the equatorial axis of the X-ray pattern was obtained by integrating the 2D pattern from 0.009 nm^−1^ on either side of the equator. The distribution of diffracted intensity along the meridional axis of the X-ray pattern (parallel to the sample axis) was obtained by integrating the 2D pattern from 0.018 nm^−1^ on either side of the meridian. Because of the arching of the meridional reflections, due to the multicellular nature of the preparations and associated variation in the orientation of myocytes^79^, to determine the spacing and the fine structure of the reflections we used the narrower integration 0.006 nm^−1^ on either side of the meridian. In a few patterns a series of very arched reflections due to connective tissue is present. These reflections have different spacings from the meridional reflections with the exception of M2 and T3 that were excluded from the analysis when affected by connective contribution. The intensity distribution of the first myosin layer line (ML1) was obtained by integrating the region between 0.030 and 0.076 nm^−1^ from the meridional axis, to exclude the contribution of the partially overlapping first actin layer line^37^.

The background intensity distribution was removed from the 1D profiles using a smoothed convex hull algorithm. The total intensities of the reflections were then obtained either by integrating the axial distribution in the corresponding regions or by a multi-gaussian fit to separate overlapping peaks. The multi-gaussian fit was applied to M3 to characterize its fine structure (fit limits 0.0676-0.0712 nm^-1^), to the M2 cluster (limits 0.042-0.049 nm^-1^) and to M5 (limits 0.115-0.118 nm^-1^).

The spacing of the reflection was determined as the intensity-weighted spacing of its component peaks. The intensity of M6 reflection was determined by integrating the axial intensity distribution between 0.137 – 0.141 nm^-1^ and its spacing was taken as the centre of gravity of the intensity distribution. This was done also for M5 when its intensity profile was too noisy for a reliable gaussian fit, within the same limits of the fit.

The intensity of the M3 reflection was corrected for the changes in the cross-meridional widths, as determined from the radial (parallel to the equatorial axis) intensity distribution in the axial regions specified above using a Gaussian fit in the region ± 0.02 nm^−1^ across the meridian. The cross-meridional width of the reflections spans about 70 pixels (FWHM), while the horizontal beam size recorded on the detector spans 18 pixels. Thus, the measured values are a good estimate of the intrinsic radial width of the reflection.

The spacing calibration was obtained taking as a reference the 14.354 nm axial spacing of the M3 reflection at rest at 27 °C ^3^.

In the experiments on the effect of temperature, the intensity of each of the analysed reflections and layer lines was first normalised for its mean value in the 27 °C patterns from each preparation and then re-normalised for the value at 35 °C. We have applied this procedure both because we collected at least one pattern at 27 °C from each of the six preparations used, allowing the same normalisation across the different preparations, and because the acquisition of more than one pattern at 27 °C in most preparations allowed an estimate of the error also for the reference intensity value.

In the experiments at different *T*_p_, to take into account the different mass of the sample crossed by the X-ray beam among different preparations and at different SL, the intensities of all the reflections examined were divided by the sum of the intensity of the equatorial reflections 1,0 and 1,1 at rest.

### Statistical analysis

Data are expressed as mean ± SD or mean ± SEM as specified. The number of preparations contributing to each protocol as well as the number n of replicates contributing to the average, which can be multiple evaluations on single trabeculae and papillary muscles, are reported in the text and in the Figure and Table legends. Statistical significance was determined using one-tailed and two-tailed *t*-test as specified and reporting the exact value of *P*.

## Data availability

All data supporting the findings of this study are available within the paper, its Supplementary Information or will be available from the corresponding authors upon reasonable request.

## Acknowledgements

We thank the European Synchrotron Radiation Facility (ESRF) for provision of synchrotron beam time, the staff of the mechanical workshop of the Department of Physics and Astronomy (University of Florence) and Jacques Gorini (ESRF) for electronic and mechanical engineering support, staff of CeSaL (University of Florence) and the staff of the Biomedical Facility (ESRF) for animal care. This project was supported by Fondazione Cassa di Risparmio di Firenze (2020.1660); European Joint Programme on Rare Diseases (IDOLS-G, EJPRD19-126); Next Generation EU programme (DM 1557 11.10.2022) in the context of the National Recovery and Resilience Plan, Investment PE8 – Project Age-It: “Ageing Well in an Ageing Society”; Italian Ministry of University and Research (MUR, PRIN2022-PNRR, P2022XPT32); and the ESRF.

## Author Contributions

Conception and design of the experiments: V.L., G.P., M.R., M.L. Investigation and data analysis I.M., M.C., M.M., M.R., G.P., I.P., P.B., T.N. Writing – Original Draft, V.L., M.R. Writing – Review & Editing, V.L., M.R., G.P., M.L. Visualization, M.C., P.B., MM. Funding Acquisition, G.P., M.L.

## Competing interests

The authors declare no competing interest

## EXTENDED DATA FIGURES AND TABLES

**Extended Data Figure 1.**
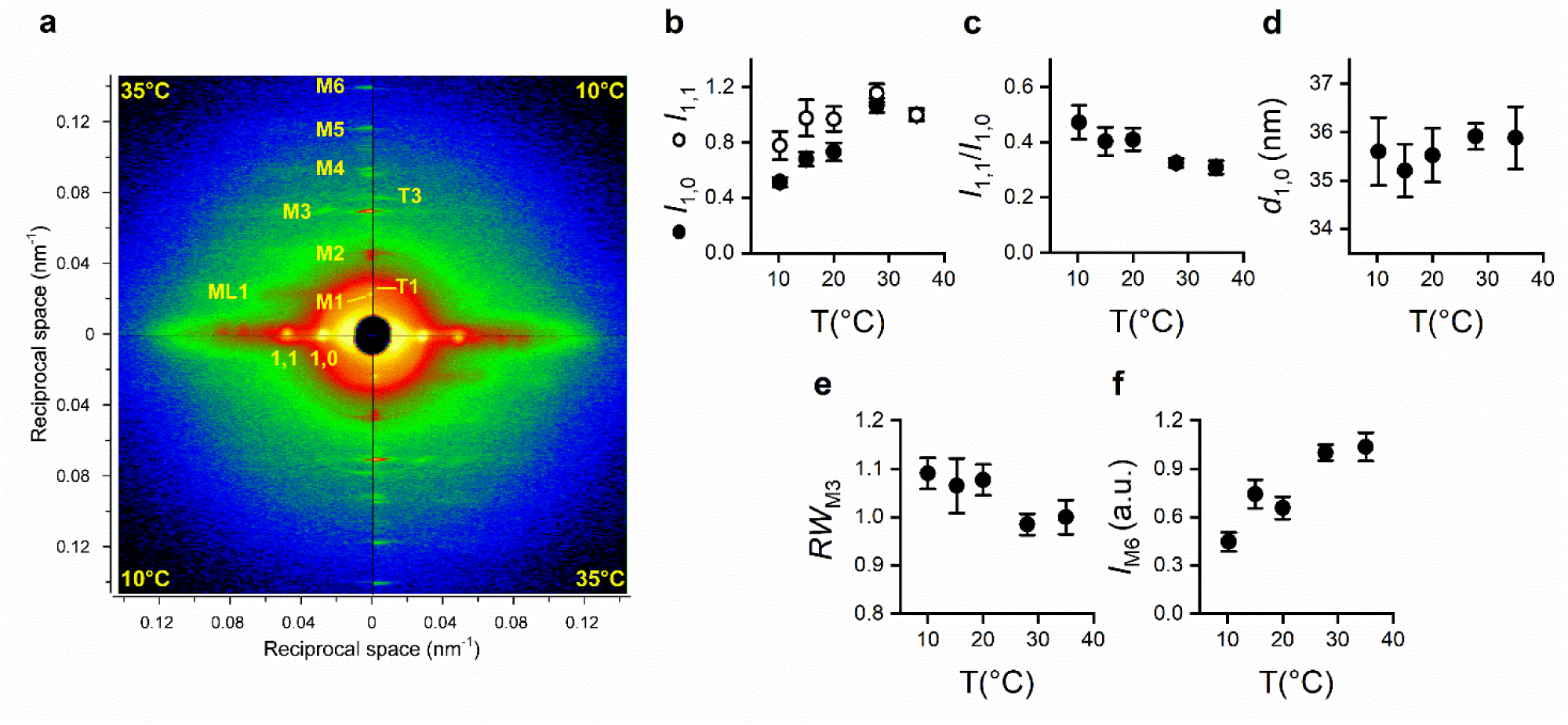
Effect of temperature on the X-ray diffraction pattern from cardiac muscle at rest. **a.** 2D-patterns collected from a papillary muscle at rest at 35 and 10 °C. The patterns have been quadrant folded (see Methods) and combined to allow direct comparison through both the meridional (vertical) and the equatorial (horizontal) axes. Upper left and lower right quadrants, 35 °C; upper right and lower left quadrants,10 °C. Each pattern has been acquired with 2×20-ms exposure windows and 1.6-m camera length. M1-M6 and T1 and T3 myosin-based and troponin-based reflections respectively. 1,0 and 1,1, strongest equatorial reflections. ML1 first myosin layer line. The arching of the intensity distributions at the level of M2 and T3 reflections, in this sample more evident in the patterns collected at 10 °C, are from connective tissue (Reconditi *et al*. 2017). **b-f.** dependence of various parameters on temperature **b.** *I*_1,0_ (filled circles) and *I*_1,1_ (open circles). **c.** *I*_1,1_ over *I*_1,0_. **d.** Spacing of the 1,0 reflection, *d*_1,0_. **e.** Radial width of the M3 reflection, *RW*_M3_, normalised for the value at 35 °C. **f.** *I*_M6_ normalised for the value a 35 °C. Data in b-f are mean ± SEM, n = 4-11, from three trabeculae and three papillary muscles.

**Extended Data Figure 2.**
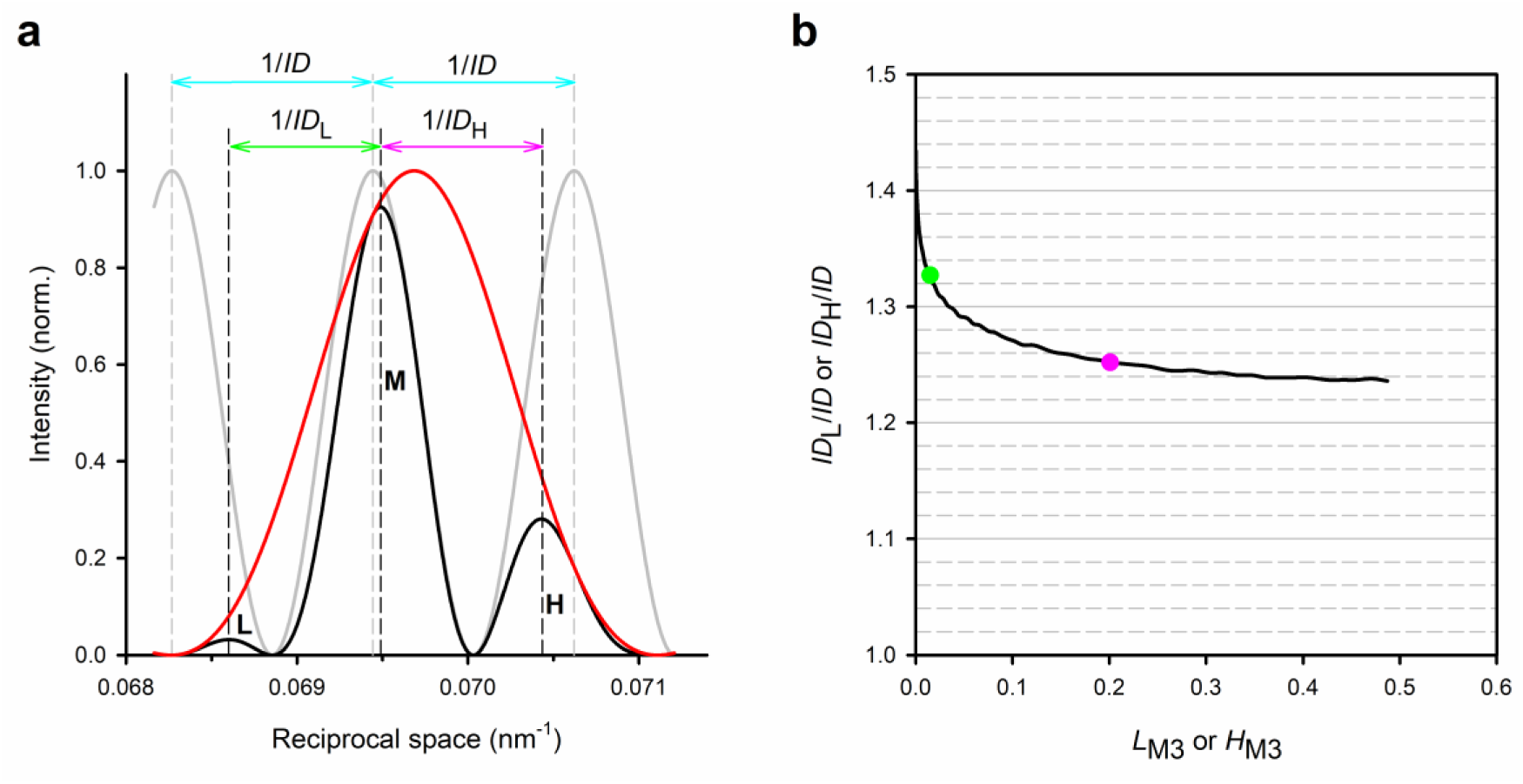
Simulation of the axial profile of the M3 intensity and relation between *ID*_L_ or *ID*_H_ and *ID*. **a.** Red, predicted intensity distribution from a single array of motors in the half-thick filament; gray, intensity distribution from two point diffractors placed at the COM of two arrays separated by a distance equal to the interference distance (*ID*); black, predicted profile of the M3 reflection from the two arrays, obtained as the product of the red and gray distributions. The dashed vertical lines indicate the positions of the peaks of the interference fringes (gray) and of the component peaks of the sampled M3 reflection (black). The horizontal arrows indicate the peak separation, from which the *ID*’s are derived as described in the text: cyan, peak separation of the interference fringes (1/*ID*); green and magenta, separation between the L and M peaks (1/*ID*_L_) and M and H peaks (1/*ID*_H_) respectively. **b.** Predicted dependence of the ratio *ID*_L_ (or *ID*_H_) over *ID* on *L*_M3_ (or *H*_M3_). The green and magenta dots refer to *ID*_L_ and *ID*_H_ respectively, measured on the intensity profiles in a.

**Extended Data Figure 3.**
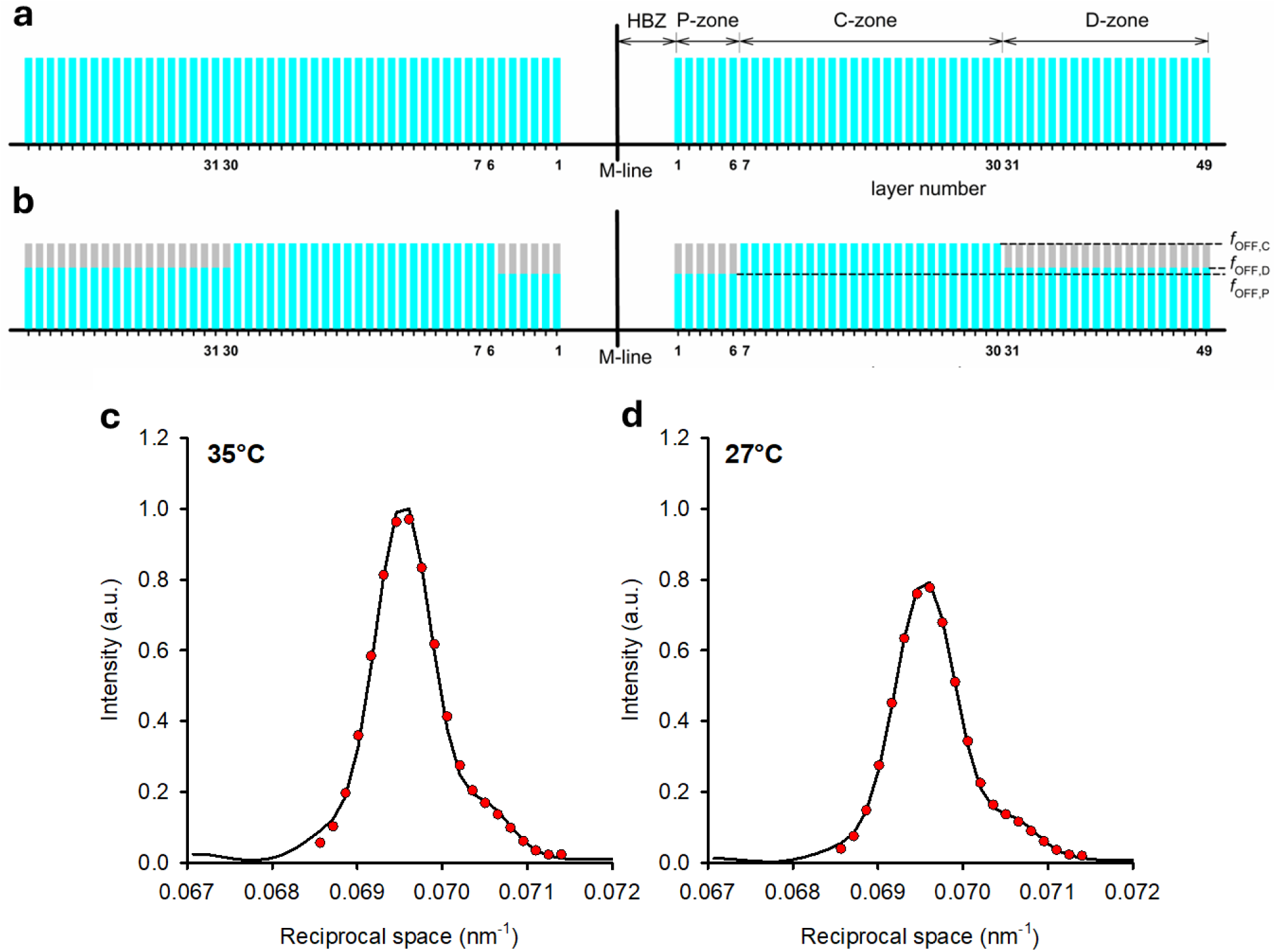
Schematic representation of the fractional occupancy of OFF and ON motor states along the thick filament and simulated M3 intensity profiles at rest at two temperatures. The layers of motors are numbered 1-49 starting from the centre of the thick filament (M-line). HBZ, axial distance of the COM of layer 1 from the centre of the filament. P-zone, layers 1-6; C-zone, layers 7-30; D-zone, layers 31-49. The length of the cyan bars is proportional to the fraction of motors in the OFF state in each zone, *f*_OFF,C_, *f*_OFF,P_, *f*_OFF,D_. The length of the gray bars represents the fraction *f*_ON_ of motors in the ON state. **a**. 35 °C: all motors are OFF. **b**. 27 °C: all motors in the C-zone are OFF, while in the P- and D-zone the fraction of OFF motors is decreased. **c.** and **d.** Observed (line) and simulated (circles) M3 intensity distribution from a papillary muscle at 35 °C (c) and 27 °C (d).

**Extended Data Figure 4.**
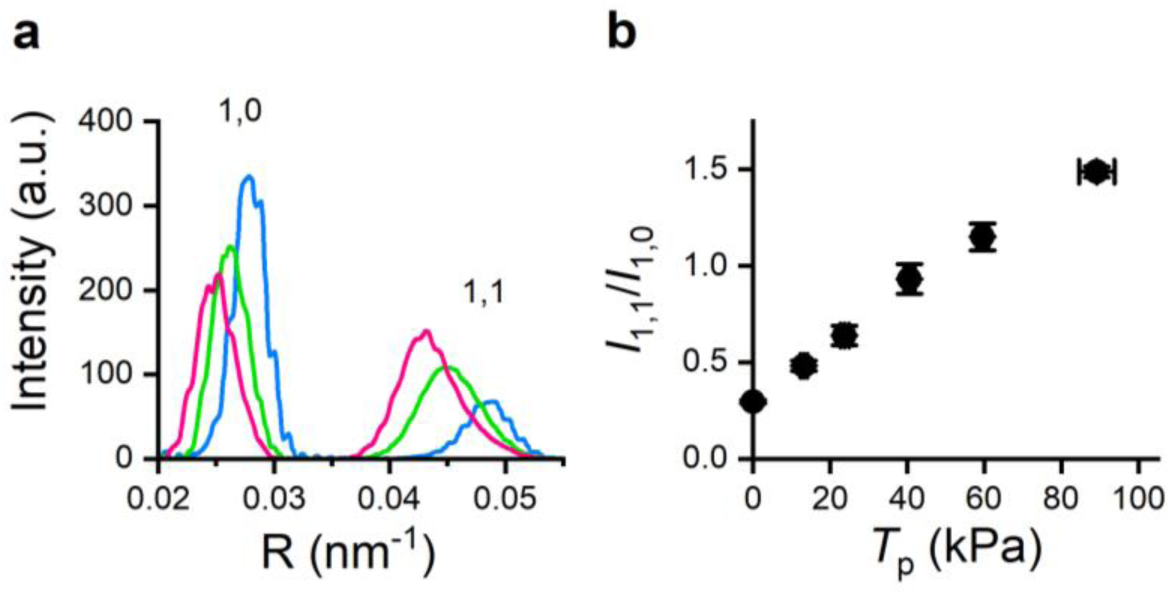
*T*_p_ dependence of the intensity of the equatorial reflections. **a.** Intensity distribution of the 1,0 and 1,1 reflections at rest (blue), at *T*_p_ 23 kPa (green) and at *T*_p_ 64 kPa (magenta). R, reciprocal space. **b.** Relation between the ratio *I*_1,1_/*I*_1,0_ and *T*_p_.

**Extended Data Figure 5.**
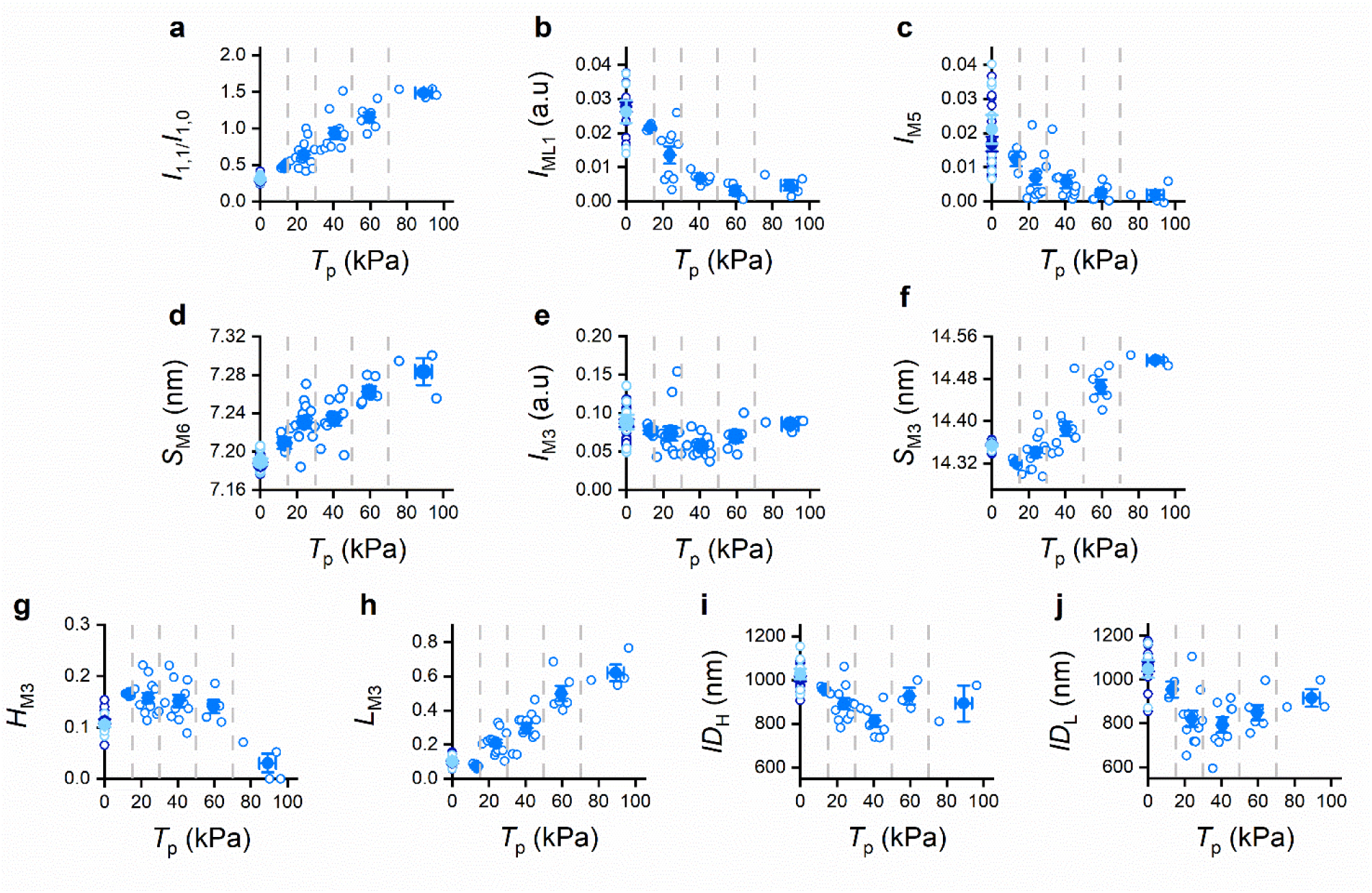
*T*_p_ dependence of the relevant X-ray parameters marking the regulatory state of the thick filament. Open circles: data collected from the different protocols as detailed in Methods; dashed gray lines, limits used for grouping pooled X-ray data in five classes of *T*_p_: 0<*T*_p_≤15; 15<*T*_p_≤30; 30<*T*_p_≤50; 50<*T*_p_≤70; *T*_p_ >70. Filled circles: mean ± SEM from the five classes. **a.** *I*_1,1_/*I*_1,0_. **b.** *I*_ML1_. **c.** *I*_M5_. **d.** *S*_M6_. **e.** *I*_M3_. **f.** *S*_M3_. **g.** *H*_M3_. **h.** *L*_M3_. **i.** *ID*_H_. **l.** *ID*_L_. The points on the ordinate are the data collected either in quiescence (blue) or in diastole (cyan). n= 3-28, from 13 preparations. CSA 130000 ± 88000 µm^2^ (mean ± SD). The mean values (± SEM) for each class are reported in Fig. 3.

**Extended Data Figure 6.**
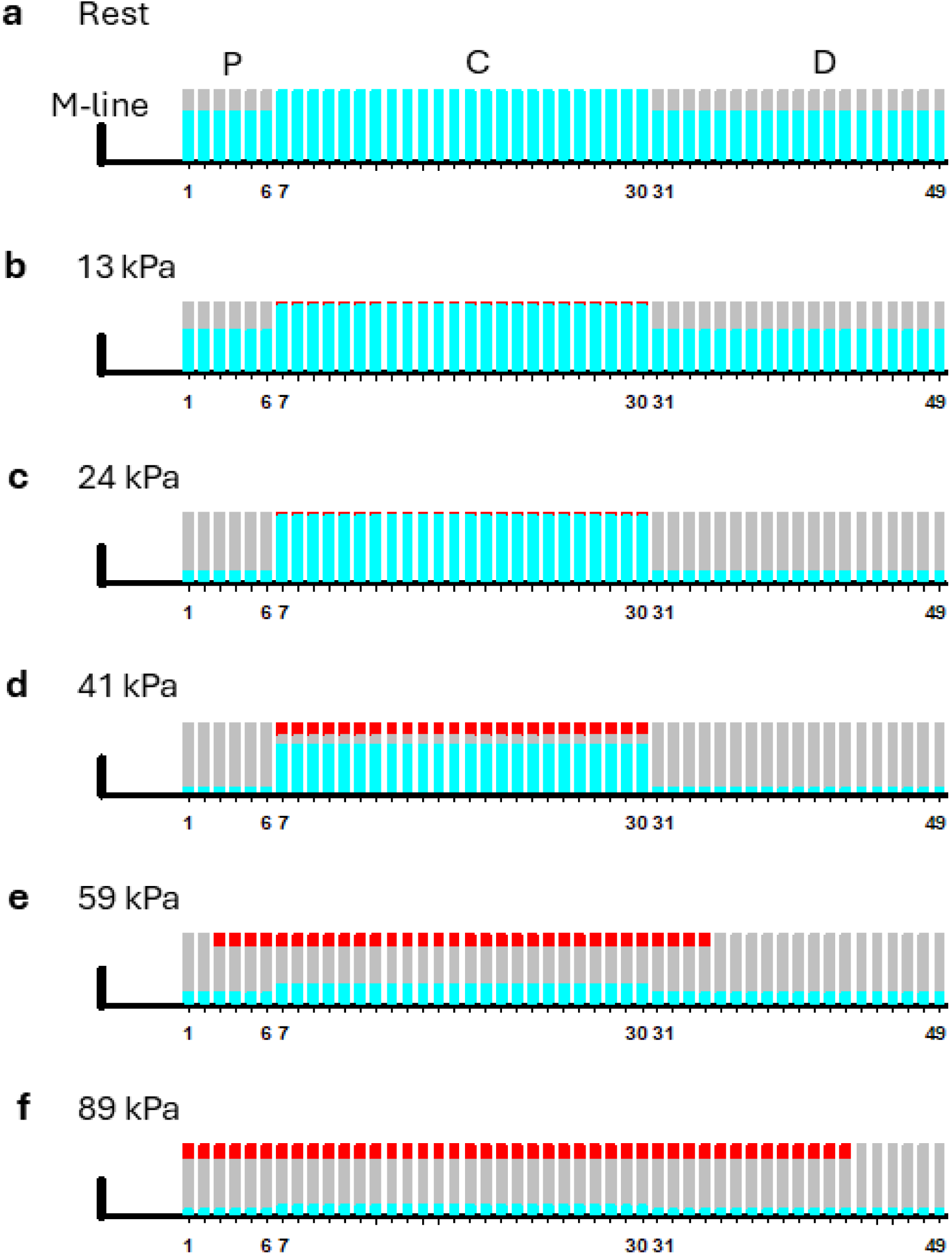
Schematic representation of motor distribution along the half-thick filament with Model 1. The length of the red bars is proportional to the fraction of attached motors. The length of the cyan bars is proportional to the fraction of OFF motors, and corresponds to the mean values found by the model simulation for *f*_OFF,C_, *f*_OFF,P_, *f*_OFF,D_. The length of the gray bars represents the fraction of ON motors. **a-f**: rest (a) and different *T*_p_ (b-f) as indicated.

**Extended Data Figure 7.**
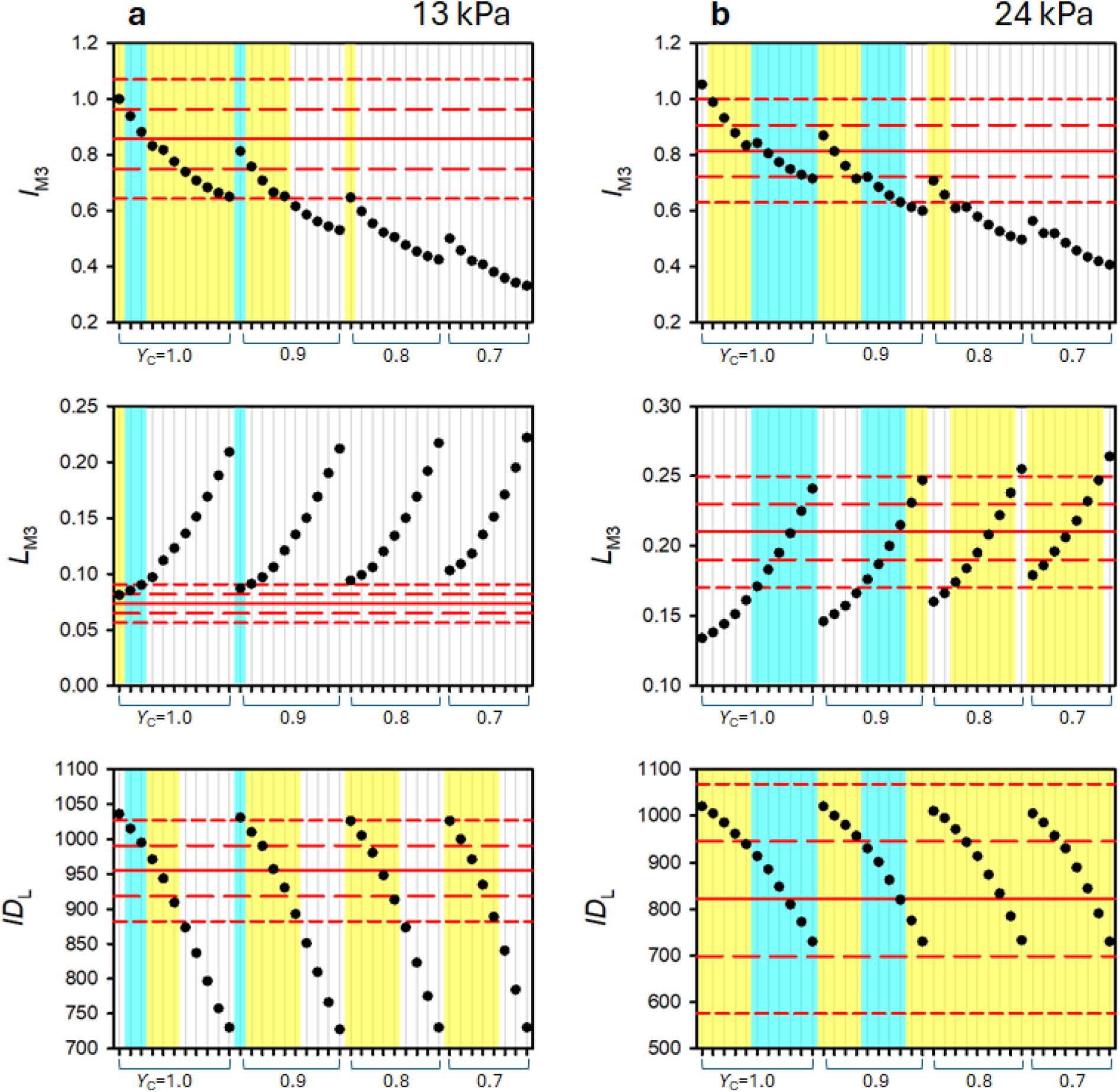
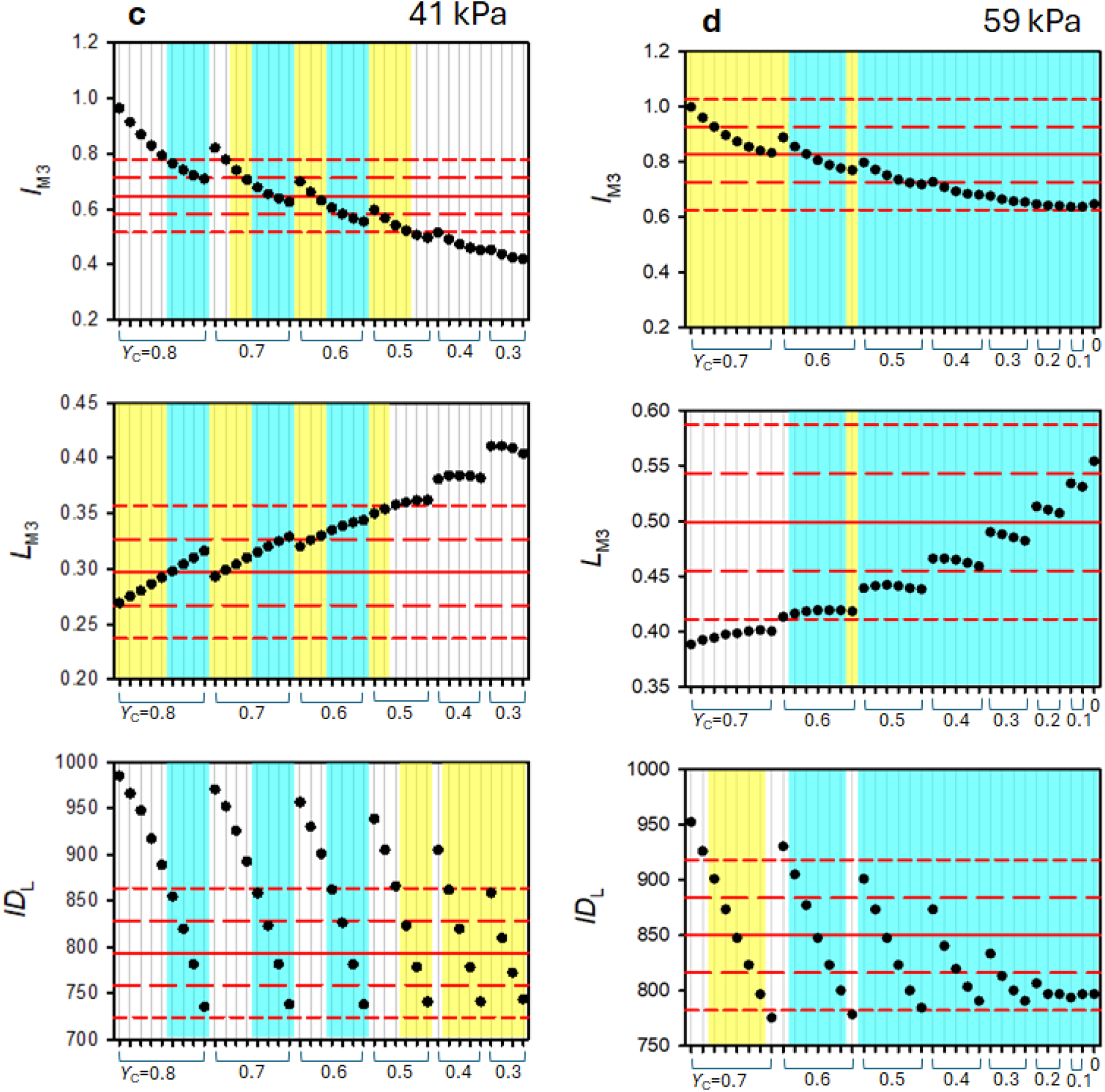
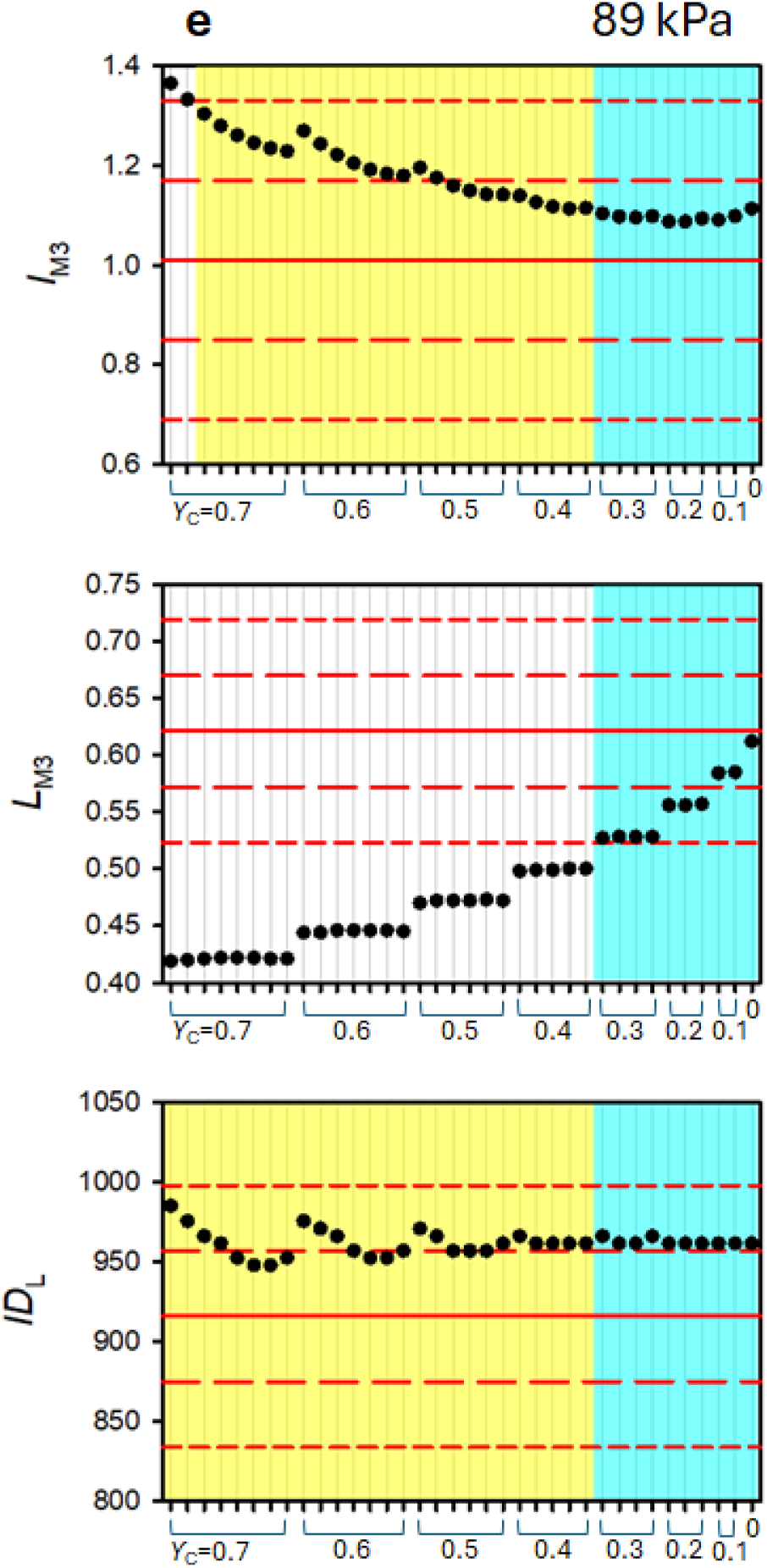
Output of Model 1 simulation for the three relevant parameters of the M3. In each panel, the red continuous line represents the observed mean value of the parameter indicated on the ordinate axis (top, *I*_M3_; middle, *L*_M3_; bottom, *ID*_L_); the red long-dashed line, mean ± SE; red short-dashed line, mean ± 2·SE. The black circles indicate the output of the model for the corresponding parameter for a given [*ϒ*_C_; *ϒ*_D_] pair, as indicated on the abscissa: for each *ϒ*_C_ value, the thicks identify the values of *ϒ*_D_ from *ϒ*_D_ = *ϒ*_C_ (left most) to 0 (right most), in steps of 0.1. The yellow shaded areas in each panel identify the [*ϒ*_C_; *ϒ*_D_] pairs for which the predicted parameter falls inside the mean ± 2·SE of the observed parameter. Cyan shaded area indicates the [*ϒ*_C_; *ϒ*_D_] pairs for which all the three parameters fall inside the mean ± 2·SE of those observed. **a-f**: different *T*_p_ values as indicated.

**Extended Data Figure 8.**
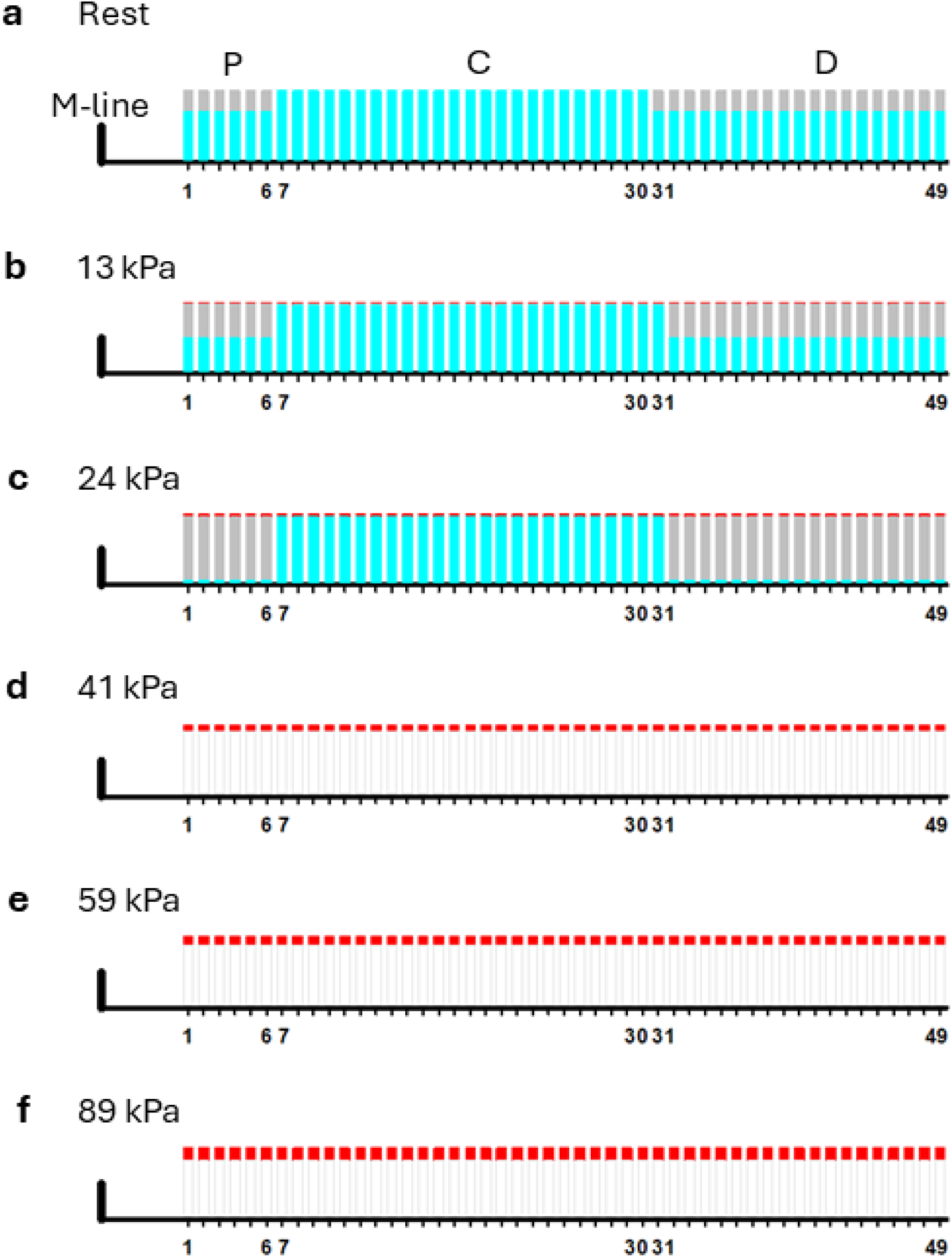
Schematic representation of motor distribution with Model 2. Layers of motors are numbered and defined as in Extended Data Fig. 5. Red, cyan and gray bars as defined in Extended Data Fig. 6. White bars indicate that the simulation fails to find a combination of *f*_OFF,C_, *f*_OFF,P_, *f*_OFF,D_ able to fit the mean values of all three parameters (*I*_M3_, *L*_M3_ and *ID*_L_). **a-f**: *T*_p_ values as

**Extended Data Figure 9.**
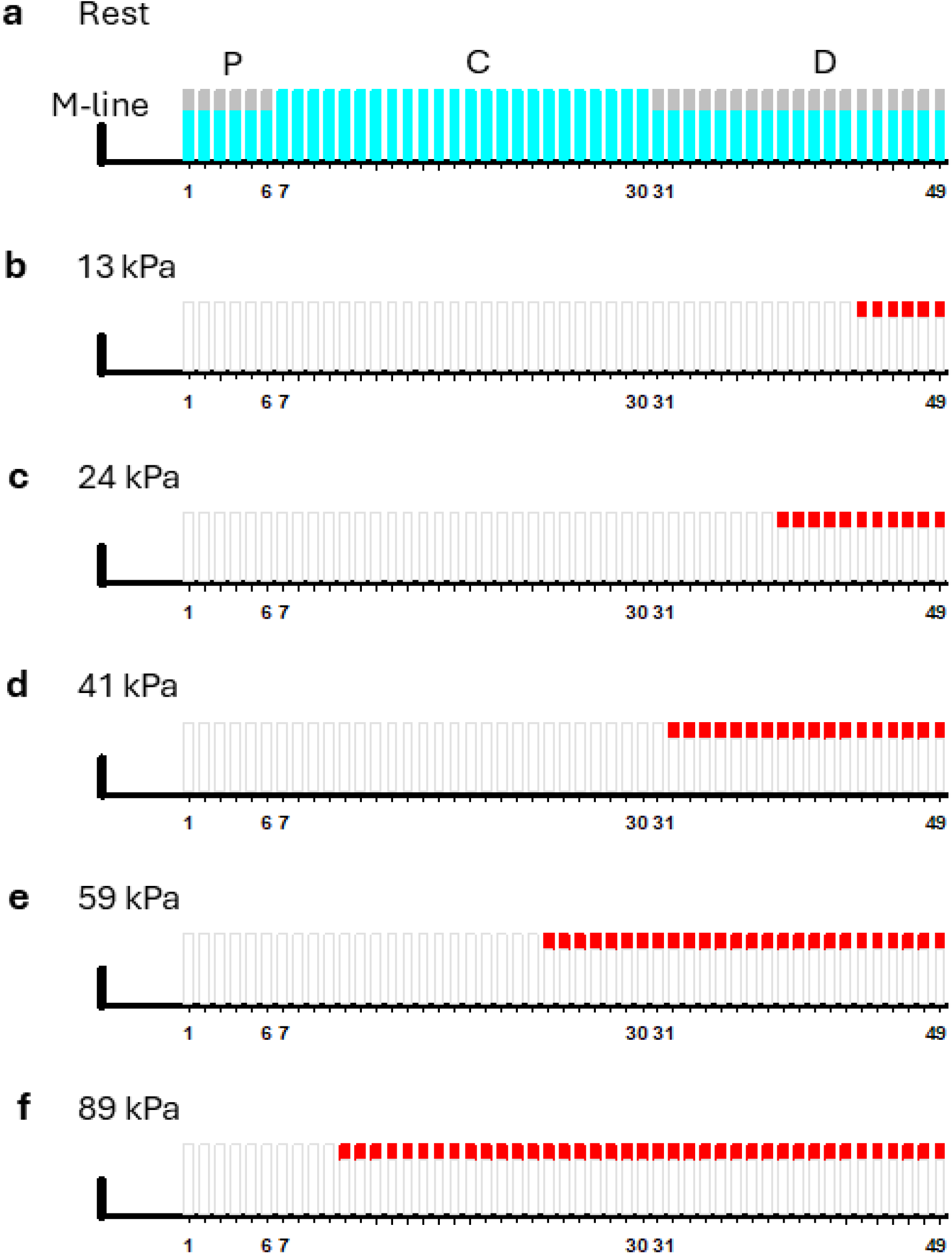
Schematic representation of motor distribution with Model 3. Layers of motors are numbered and defined as in Extended Data Fig. 5. Red, cyan and gray bars as defined in Extended Data Fig 6. White bars indicate that the simulation fails to find a combination of *f*_OFF,C_, *f*_OFF,P_, *f*_OFF,D_ fitting the mean values of all three parameters (*I*_M3_, *L*_M3_ and *ID*_L_). **a-f**: *T*_p_ values as

**Extended Data Figure 10.**
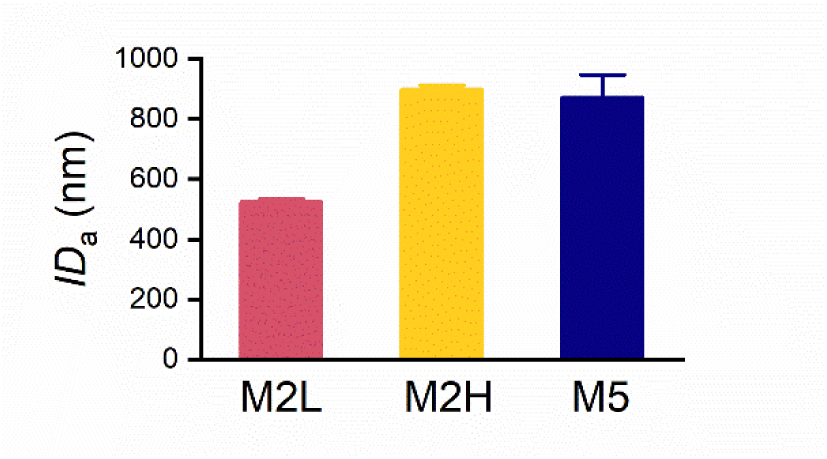
Interference distances estimated from the fine structure of the M2 and M5 reflections. Interference distance (*ID*_a_) from intensity profiles as in Fig. 2b (M2) and c (M5) at temperature 35 °C. *ID*_a_ is calculated as the reciprocal of the peak separation of the two low angle component peaks (M2L, red) and of the two high angle component peaks (M2H, yellow) and as the reciprocal of the peak separation of the three component peaks of M5 (blue). The corresponding spacings are 22.71 ± 0.06 nm (M2L); 21.40 ± 0.01 nm (M2H) and 8.603 ± 0.007 nm (M5) (mean ± SEM, n=4). Thus M2H and M5 are indexed as the 2^nd^ and the 5^th^ order of a fundamental ∼43 nm periodicity, while M2L is the 2^nd^ order of a fundamental ∼45 nm periodicity. M2H (yellow) and M5 (blue) are characterized by an *ID*_a_ (898 ± 14 nm and 871 ± 76 nm respectively) that is much larger than that of M2L (525 ± 9 nm), indicating that the structure that is responsible for the axial perturbation with fundamental 43 nm periodicity is distributed through nearly the whole thick filament, while the structure responsible for the axial perturbation with fundamental 45 nm periodicity is limited to a more central zone of the thick filament.

**Extended Data Table 1.** Values of relevant parameters of M3 and M6 at rest and at *T*_p_ = 13 kPa. Fractional contributions to the intensity of the M3 reflection of the low angle peak, *L*_M3_, and of the high angle peak, *H*_M3_. Spacings of the M3, *S*_M3_, and M6, *S*_M6_. Data are mean ± SEM. The statistical significance of the differences between rest and *T*_p_ = 13 kPa is expressed by *P* estimated with one-tailed *t*-test. *df*, degree of freedom.

|  | Rest | 13 kPa | *df* | *P* |
| --- | --- | --- | --- | --- |
| $L_{M3}$ | $0.099 \pm 0.006$ | $0.073 \pm 0.009$ | 29 | 0.011 |
| $H_{M3}$ | $0.110 \pm 0.003$ | $0.1654 \pm 0.0023$ | 29 | $3 \cdot 10^{-15}$ |
| $S_{M3}$ (nm) | $14.3548 \pm 0.0012$ | $14.322 \pm 0.004$ | 29 | $6 \cdot 10^{-9}$ |
| $S_{M6}$ (nm) | $7.1880 \pm 0.0014$ | $7.209 \pm 0.006$ | 29 | 0.001 |

## Notes

### Competing Interest Statement

The authors have declared no competing interest.

### Summary of Updates

This version of the manuscript has been revised to update Extended Data Figure 10.

