## Supplementary Note 1,2 and 3 and Supplementary Table 1, 2 and 3 for "An integrated picture of the structural pathways controlling the heart performance"

### Supplementary information

#### Supplementary Note 1

##### Estimate of the interference distance

The fine structure of the M3 reflection along the meridional axis is governed by the X-ray interference between the two arrays of motors on the two halves of the thick filament that samples the reflection with peaks of different intensity (L, low angle peak; M, medium angle; H, high angle, Extended Data Fig. 2a). The interference distance *ID*, i.e. the distance between the centres of mass (COM) of the two motor arrays, can be derived from the positions $R_i$ and $R_{i+1}$ in the reciprocal space of two consecutive interference peaks according to the formula:

$$\boldsymbol{ID} = \frac{\mathbf{1}}{\boldsymbol{R}_{\mathbf{i+1}} - \boldsymbol{R}_{\mathbf{i}}}$$

However, the observed intensity profile is the product of the profile generated by the ~14.5 nm periodicity of the array of heads in each half-thick filament and the interference fringes generated by the two motor arrays in each thick filament that act as two point-diffractors placed at the array COM The result is that the positions of the interference peaks are displaced towards each other (Huxley *et al.*, 2006), thus the interference distance deduced from their positions ($ID_L$ or $ID_H$, depending on whether the M-L separation or the M-H separation is used) is somewhat larger (by ~20%) than the true distance between the COM of the two motor arrays (Extended Data Fig. 2b).

Moreover, due to the asymmetry of the fine structure, *ID* calculated from the positions in the reciprocal space of either the L ($R_L$) and M ($R_M$) peaks ($ID_L$) or the M and H ($R_H$) peaks ($ID_H$) may be slightly different.

$$ID_{\mathrm{L}} = \frac{\mathbf{1}}{\boldsymbol{R}_{\mathbf{L}} - \boldsymbol{R}_{\mathbf{M}}} \neq ID_{\mathrm{H}} = \frac{\mathbf{1}}{\boldsymbol{R}_{\mathbf{M}} - \boldsymbol{R}_{\mathbf{H}}}$$

The relation between the $ID$ and $ID_L$ or $ID_H$ shown in Extended Data Fig. 2b is derived assuming an array of motors where all the 49 layers on each half thick filament contribute equally to M3. In estimating $ID$, it is preferable to use whichever of L or H peak has the higher intensity. This is because both the S/N is better for the higher intensity peak and the relation shown in Extended Data Fig. 2b is less steep for higher values of either $L_{M3}$ or $H_{M3}$. Also for this reason, to find the best fit in modelling the M3 profile at rest at different temperatures we privileged the H peak, while for modelling the M3 parameters at different $T_P$ we privileged the L peak.

#### Supplementary Note 2

##### Structural model with different regional sensitivity to cooling

The observed changes in the intensity and fine structure of the M3 reflection due to the perturbing action of cooling on the OFF motor conformation have been simulated with a simple structural model of the sarcomere. Since we are modelling a single reflection with a small spacing change (<0.25%), each layer of six myosin motors (three dimers) can be represented as a point diffractor (Linari *et al*. 2000). In the model, two arrays of equally spaced point diffractors represent the layers of myosin motors along each half thick filament (Extended Data Fig. 3a,b). The position of each point diffractor coincides with the axial COM of the corresponding motor layer (Reconditi 2006). Each array is divided in three segments or zones: proximal (P), central (C) and distal (D) zone with 6, 24 and 19 diffractors respectively (Luther *et al*. 2008). The different fractions of motors in the OFF conformation are reflected in a different weight $f$ of the diffracting power of the point diffractors in the three regions. This is equivalent to the assumption that, within each zone and through the many thick filaments across a sarcomere, the motors maintaining the OFF conformation and those leaving it are randomly distributed. The extent of the bare zone (BZ) between the two arrays is taken here as the axial distance between the proximal point diffractors in the two arrays. Note that the extent of the BZ normally indicates the separation between the rod-head junction of the proximal motors in the two arrays (Reconditi *et al*. 2011), while here it indicates the separation of their COM The calculated profile is then convoluted with a gaussian function that represents the shape of the X-ray beam as recorded directly on the detector. It must be noted, however, that the meridional axis of the diffraction pattern may be somewhat rotated relative to the detector axes, along which the shape of the beam is determined. Thus, the shape of the beam profile along the meridional axis of the pattern depends on the pattern orientation on the detector and is always larger than what measured in the vertical direction.

The parameters to be estimated to fit the observed profiles are:

$d_0$: the periodicity of the diffractors along each array

HBZ (Half Bare Zone): the distance from the COM of the proximal point diffractor from the centre of the sarcomere

$f_{OFF,P}$: the fraction of OFF motors in the P-zone

$f_{OFF,C}$: the fraction of OFF motors in the C-zone

$f_{OFF,D}$: the fraction of OFF motors in the D-zone

$\sigma_{beam}$: the effective size of the (gaussian) beam profile along the meridional axis

$\alpha$: a scaling factor to fit the overall intensity of the calculated profile to that observed.

In the model, the total intensity of M3 scales as the square of the fraction $f$ of contributing motors (thus excluding the disordered motors; Linari *et al.* 2000), and as the scaling factor α, which is constant independent of temperature.

The profile of M3 along the meridional axis is calculated at each value of the reciprocal space coordinate $R$ of the detector pixels recording the intensity of the reflection:

$$\mathrm{I}(R) = \left( \sum_{i=1}^{6} f_{\mathrm{OFF,P}} \cdot \cos\{2\pi \cdot [\mathrm{HBZ} + (i-1) \cdot d_0] \cdot R\} + \sum_{i=7}^{30} f_{\mathrm{OFF,C}} \cdot \cos\{2\pi \cdot [\mathrm{HBZ} + (i-1) \cdot d_0] \cdot R\} + \sum_{i=31}^{49} f_{\mathrm{OFF,D}} \cdot \cos\{2\pi \cdot [\mathrm{HBZ} + (i-1) \cdot d_0] \cdot R\} \right)^2$$

For the simulation of the observed M3 intensity distribution along the meridional axis, this intensity profile is convoluted with a gaussian beam profile of width $\sigma_{beam}$ and then multiplied by the scaling factor α.

The best fit to the data is obtained searching the values of the parameters that minimize the sum of the squared difference between the intensity output of the model and that measured at each position (least-squares curve fitting). The model has been implemented in Mathcad 15 (PTC Inc.), which uses a Levenberg-Marquardt algorithm to find the minimum sum of the squared residuals.

The first step in the modelling is aimed at determining the value of HBZ, which is then assumed not to be influenced by the changes in temperature (Reconditi *et al.* 2011) and consequently set as a fixed parameter for the fit on the M3 intensity profiles at the different temperatures. For this, we have fitted the profile at 35 °C with the assumption that at this temperature all the motors are in the OFF configuration throughout the three regions, i.e. $f_{OFF,P} = f_{OFF,C} = f_{OFF,D} = 1$ (Caremani *et al.* 2019; Ovejero *et al.* 2022). The model has been applied to the profile of M3 obtained by adding the patterns at 35 °C from four preparations, to improve the signal-to-noise ratio of the intensity distribution and minimize the error for the evaluation of HBZ. The value found for HBZ is 79.8 nm. Together with HBZ, which characterises the fine structure of the reflection, the fit also determines the constant scaling factor α. In principle, the apparent beam size $\sigma_{beam}$ could vary, by a small amount, among preparations and among exposures of different regions of each preparation. However, we found that the results of the fit are more consistent if we also fix this value, which from the added patterns at 35 °C is $2.96 \cdot 10^{-4}$ $nm^{-1}$, 30% larger than the $2.28 \cdot 10^{-4}$ $nm^{-1}$ of the vertical beam size measured on the detector, for the reason explained above. Then, keeping the values of HBZ, $\sigma_{beam}$ and the intensity scaling factor $\alpha$ fixed, the model simulation has been run to fit the M3 intensity profiles of each preparation at the different temperatures with the fractions $f_{OFF,P}$, $f_{OFF,C}$ and $f_{OFF,D}$ as free parameters. The results of the fit are shown in Fig. 2k. The graphic representation of the fractional occupancy of either the OFF or the ON state for each crown at 27 °C is shown in Extended Data Fig. 3b ($f_{OFF}$, cyan; $f_{ON}$, gray). In Extended Data Fig. 3c,d the M3 intensity distribution obtained from the fit of the model is superimposed on that observed for 35 and 27°C.

The results of the model simulation indicate that $f_{OFF,P}$ (Fig. 2k, green) and $f_{OFF,D}$ (Fig. 2k, brown) are reduced in the same way by lowering temperature and much more than $f_{OFF,C}$. Thus, the disordering effect of cooling is equally effective in the two zones in which MyBP-C is absent. For all the temperatures, the model output indicates that $f_{\mathrm{OFF,P}}$ and $f_{\mathrm{OFF,D}}$ are not statistically different. The two-tailed $t$ test provides the following $P$ values: 35°C, $P = 0.47$; 27°C, $P = 0.72$; 20°C, $P = 0.48$; 15°C, $P = 0.18$; 10°C, $P = 0.15$; $n = 4$, 6 degrees of freedom.

Taking $f_{\mathrm{OFF,P}} = f_{\mathrm{OFF,D}}$, their common value can be calculated as the weighted mean of the results of the fit. At 27 °Cat rest, the values for the fraction of motors in the OFF conformation in P, C and D are 0.70 ± 0.17 (green circle at 27 °C in Fig. 2k), 1.07 ± 0.05 (black circle) and 0.77 ± 0.06 (brown circle) respectively (mean ± SE). The mean $\overline{x_p}$ of the two values for $f_{\mathrm{OFF,P}}$ and $f_{\mathrm{OFF,D}}$ weighted for their errors and the corresponding error $\overline{\sigma_p}$ are given by:

$$\overline{x_p} = \Sigma \frac{\overline{x_i}}{\sigma_i^2} / \Sigma \frac{1}{\sigma_i^2}, \; \overline{\sigma_p} = 1 / \sqrt{\Sigma \frac{1}{\sigma_i^2}}$$

With $\overline{x_1} = 0.70, \overline{x_2} = 0.77, \sigma_1 = 0.17, \sigma_2 = 0.06$ we found $f_{\mathrm{OFF,P}} \equiv f_{\mathrm{OFF,D}} = 0.76 \pm 0.06$. Following the normalization for $f_{\mathrm{OFF,C}} = 1.07$, the new set of data at rest at 27 °C, graphically represented in Fig. 2m, are: $f_{\mathrm{OFF,C}} = 1$, $f_{\mathrm{OFF,P}} \equiv f_{\mathrm{OFF,D}} = 0.71$. The new estimate of HBZ for the best fit of the M3 parameters at rest at 27°C is 79.4 nm, consistent with the 79.8 nm found before averaging $f_{\mathrm{OFF,P}}$ and $f_{\mathrm{OFF,D}}$.

#### Supplementary Note 3

##### Structural model for different systolic forces

The structural model used to simulate the effect of lowering temperature on M3 intensity distribution at rest and to predict the fraction and regional localization of the motors leaving the OFF state has been implemented to simulate the M3 intensity profiles at different $T_p$ by introducing the contribution of the attached motors and their detached partner in the dimer to the reflection (Piazzesi *et al.* 2002; Brunello *et al.* 2007). As a starting point, the model simulation uses the fraction of motors in the OFF conformation in P, C and D zones predicted by the model simulation at rest at 27 °C (Fig. 2m, reported in Fig. 5a, rest): $f_{OFF,C} = 1, f_{OFF,P} \equiv f_{OFF,D} = 0.71$ (see Supplementary Note 2). The COM of the attached motors differs from that of the dimer in the OFF state by 10.5 nm (Reconditi *et al.* 2011; 2017). Moreover, the diffracting power $w$ of an attached dimer (attached motor and detached partner) is 4.8 the mean diffracting power of a motor in the OFF conformation (Reconditi *et al.* 2017). We indicate with $f_A$ the fraction of attached motors in each layer of 6 motors that contribute to M3. With $n$ contributing layers in the half thick filament, the total number $N_A$ of attached motors in each half thick filament is $n \cdot 6 \cdot f_A$. With the protocols used to vary $T_p$, (See Methods, Caremani *et al.* 2016) $N_A$ scales linearly with the force $T_p$. The maximal systolic force $T_{p,max}$ is taken as 110 kPa (Schouten *et al.* 1990; Reconditi et al., 2017), at which $n = 49, f_A = 0.22$ (all the motor layers contribute equally), and $N_A = 65$ (Reconditi *et al.* 2017).

At any force the attached motors contribute from layers between $n_i$ and $n_e$, 1 and 49 being respectively the proximal and distal layers relative to the M-line (Brunello *et al.* 2020).

The intensity profile of the M3 reflection along the meridional axis as a function of the reciprocal space coordinate $R$ is then calculated as:

$$\mathrm{I}(R) = \left( \sum_{i=1}^{6} f_{\mathrm{OFF,P}} \cdot \cos\{2\pi \cdot [HBZ + (i-1) \cdot d_0] \cdot R\} + \right.$$

$$+ \sum_{i=7}^{30} f_{\mathrm{OFF,C}} \cdot \cos\{2\pi \cdot [\mathrm{HBZ} + (i-1) \cdot d_0] \cdot R\} +$$

$$+ \sum_{i=31}^{49} f_{\mathrm{OFF,D}} \cdot \cos\{2\pi \cdot [\mathrm{HBZ} + (i-1) \cdot d_0] \cdot R\}$$

$$\left. + \sum_{i=n_i}^{n_e} f_{\mathrm{A}} \cdot w \cdot \cos\{2\pi \cdot [\mathrm{HBZ} + \Delta z + (i-1) \cdot d_0] \cdot R\} \right)^2$$

The general constraint for model simulation is that the factor $f_A \cdot (n_e - n_i + 1)$ (that is $N_A/6$) scales in direct proportion with $T_p$. The strategy we used to simulate the data at different temperatures was to find the best fit of the intensity distributions of the M3 reflection (Extended Data Fig. 3c,d). However, the M3 intensity profiles in systole for the same class of $T_p$ were less homogeneous than at rest for the same temperature. Thus, differently from the simulation of temperature effect at rest, in looking for the best structural model of any class of $T_p$, we took the option to simulate the relevant parameters that are extracted from the M3 intensity distribution, in particular: $I_{M3}$, $S_{M3}$, $L_{M3}$ and $ID_L$ (the separation between M and L peaks used to estimate $ID$).

For any given $T_p$ (first column in Supplementary Tables 1-3) and number of attached motors (second column) the model output is calculated for pairs of fractional reduction of $f_{OFF,C}$, $f_{OFF,D}$ values (with $f_{OFF,P}$ the same as $f_{OFF,D}$). Indicating with $\Upsilon_C$ and $\Upsilon_D$ respectively the values of $f_{OFF,C}$ and $f_{OFF,D}$ relative to their values at rest (that is $f_{OFF,C} = \Upsilon_C$ and $f_{OFF,D} = 0.71 \cdot \Upsilon_C$), we changed $\Upsilon_C$ and $\Upsilon_D$ in steps of 0.1, in the range $0 \leq \Upsilon_C \leq 1.0$ and $0 \leq \Upsilon_D \leq \Upsilon_C$, adjusting $d_0$ to get the best fit to the observed mean value of $S_{M3}$. A [$\Upsilon_C$; $\Upsilon_D$] pair is considered compatible with the data when the model output with that pair gives an estimate of the other three parameters ($I_{M3}$, $L_{M3}$, $ID_L$) that falls inside the range of ±2·SEM ($p<0.05$) from the observed mean values.

The model was tested for three different hypotheses on the localization of attached motors and their progression with force: the first hypothesis (Model 1, Extended Data Fig. 6 and 7 and Supplementary Table 1) is that motors attach first in the C-zone and further attachments with increase in $T_p$ progress towards the periphery of the thick filament; the second hypothesis (Model 2, Extended Data Fig. 8) is that motors attach uniformly throughout the thick filament ($n_i = 1$, $n_e = 49$ for all $T_p$) and increase in $T_p$ is achieved with homogeneous increase in the fraction of attached motors throughout the filament, i.e. $f_A$ increases linearly with $T_p$ (Supplementary Table 2); the third hypothesis (Model 3, Extended Data Fig. 9) is that motors attach first at the edges of the thick filament and then progress towards the centre of the thick filament with the increase in $T_p$, with $f_A = 0.22$ and $n_e = 49$ for all $T_p$, and the length of the array of attached motors (represented by $n_e$-$n_i$) increasing linearly with $T_p$ (Supplementary Table 3).

*Model 1 output*

Model 1 implies that the first motor attachments occur uniformly in the C-zone of the thick filament ($n_i = 7$ and $n_e = 30$) and $f_A$ increases linearly with force until, at 41 kPa, it attains the value of 0.167 (third column in Supplementary Table 1), at which there is one attached motor per layer (0.167·6). This condition is adequate for the cooperative thin filament activation by attached motors to allow further attachments to spread symmetrically out of the C-zone. Thus for $T_p$ from 13 to 41 kPa, $N_A$ rises from 8 to 24 (second column in Supplementary Table 1) and the fraction $f_A$ rises from 0.053 to 0.167 (red bar in Fig. 5, 2$^{nd}$ and 3$^{rd}$ panel, and in Extended Data Fig.6, b-d). At forces larger than 41 kPa the increase of $f_A$, $\Delta f_A$ (= $f_A$ - 0.167), is directly proportional to the increase of force $\Delta T_p$ (=$T_p$ - 41 kPa). $n_i$ and $n_e$ must satisfy the relation ($n_e$ - $n_i$ +1) = (49·0.22/110 kPa) ·$T_p$ / $f_A$. At $T_p$ 59 kPa: $f_A$ =0.181 and $n_i$ =3, $n_e$ = 34 (Extended Data Fig. 6e). Once the symmetrical spread of attachments reaches layer 1 and saturates the P-zone, attachments progress only through the distal region of the D-zone. At $T_p$ 89 kPa: $f_A$ =0.204 and $n_i$ =1, $n_e$ = 43 (Fig 5, 4th panel, and Extended Data Fig.6f).

In Supplementary Table 1 are reported the values of $f_A$ (third column), $n_i$ (fourth column), and $n_e$ (fifth column) for all $T_p$ values. $n_e$ is predicted to reach layer 49 at 110 kPa, where $f_A$ = 0.22.

A detailed description of the dependence of the three parameters $I_{M3}$, $L_{M3}$ and $ID_L$ on [$\Upsilon_C$; $\Upsilon_D$] for Model 1 is given in Extended Data Fig. 7. The yellow area in each plot identifies the values of [$\Upsilon_C$; $\Upsilon_D$] pairs for which the corresponding parameter falls inside the range of ±2·SEM (short dashed red lines) ($p<0.05$) from the observed mean value (continuous red lines). The cyan area identifies the values of [$\Upsilon_C$; $\Upsilon_D$] pairs for which the above condition is satisfied by all three parameters. The corresponding $f_{OFF,C}$ (=$\Upsilon_C$) and $f_{OFF,D}$ (=0.71·$\Upsilon_C$) are reported in Supplementary Table 1, 6th and 7th column respectively. The errors associated to the mean values represent the extreme limits that can be assumed by $f_{OFF}$ in C and D-zone still able to fit all three parameters in the range ±2·SEM. The power of Model 1 for interpreting the changes in the parameters of the M3 intensity distribution becomes strikingly evident in the plots concerning $T_p$ at 41 kPa: the cyan shaded area, which represents the ensemble of [$\Upsilon_C$; $\Upsilon_D$] pairs that fit the mean values of all three parameters, appears dispersed in three clusters of pairs. Actually, the [$\Upsilon_C$; $\Upsilon_D$] pairs point to a quite restricted low fraction of OFF motors in the D-zone and a quite restricted high fraction of OFF motors in the C-zone. In fact, 41 kPa is the $T_p$ value at which the D-zone has already lost most of OFF motors, which are still largely present in the C-zone. Comparing the yellow dashed areas in the plots of the three parameters at 41 kPa we can see that such a sharp conclusion is determined by $ID_L$, the parameter for which yellow and cyan areas coincide.

*Model 2 output*

In Model 2, motors attach uniformly throughout the thick filament ($n_i$ = 1, $n_e$ = 49 for all $T_p$) and increase in $T_p$ is achieved with homogeneous increase in the fraction of attached motors throughout the filament, i.e. $f_A$ increases linearly with $T_p$ (Extended Data Fig. 8 and Supplementary Table 2). In the simulation with Model 2, pairs of [$\Upsilon_C$; $\Upsilon_D$] values that can fit the observed mean values of all three parametes ($I_{M3}$, $L_{M3}$ and $ID_L$) exist only for the lower $T_p$ (13 and 24 kPa). For higher $T_p$ (41, 59 and 89 kPa) the simulation with Model 2 fails to give any [$\Upsilon_C$; $\Upsilon_D$] pair fitting the observed mean values of all three parameters.

*Model 3 output*

In Model 3 motors attach first at the edges of the thick filament and then progress towards the centre of the thick filament with the increase in $T_p$. $f_A$ (= 0.22) and $n_e$ (= 49) are the same for all $T_p$, and the length of the array of attached motors (represented by $n_e$-$n_i$) increases linearly with $T_p$ (Extended Data Fig. 9 and Supplementary Table 3). The simulation demonstrates that for all $T_p$, no [$\Upsilon_C$; $\Upsilon_D$] pair exists that can fit the observed mean values for all three parameters under the assumption in Model 3.

### Supplementary Tables

| $T_p$ (kPa) | $N_A$ | $f_A$ | $n_i$ | $n_e$ | $f_{OFF,C}$ | $f_{OFF,D}$ |
|---|---|---|---|---|---|---|
| 0 (Rest) | 0 | 0 | - | - | 1.00 | 0.71 |
| 13 | 8 | 0.053 | 7 | 30 | 0.95 ± 0.05 | 0.60 ± 0.04 |
| 24 | 14 | 0.098 | 7 | 30 | 0.95 ± 0.05 | 0.18 ± 0.18 |
| 41 | 24 | 0.167 | 7 | 30 | 0.70 ± 0.10 | 0.10 ± 0.10 |
| 59 | 35 | 0.181 | 3 | 34 | 0.30 ± 0.30 | 0.18 ± 0.18 |
| 89 | 53 | 0.204 | 1 | 43 | 0.15 ± 0.15 | 0.10 ± 0.10 |

**Supplementay Table 1. Parameters used for the simulation of Model 1 and results from the fit.** Columns from the left: $T_p$; $N_A$, number of attached motors, which scales linearly with $T_p$; $f_A$: fraction of attached motors in each layer of 6 motors; $n_i$ and $n_e$: first and last motor layer of the array with attached motors. $f_{OFF,C}$ and $f_{OFF,D}$: fraction of OFF motors in the C-zone and in the D- (and P-) zone respectively; mean values and range for the fit of the observed parameters as described in the text. At $T_{p,max}$ (=110 kPa), $N_A$= 65, $f_A$= 0.22, $n_i$= 1, $n_e$= 49, $f_{OFF,C}$= $f_{OFF,D}$= 0.

| $T_p$ (kPa) | $N_A$ | $f_A$ | $n_i$ | $n_e$ | $f_{OFF,C}$ | $f_{OFF,D}$ |
|---|---|---|---|---|---|---|
| 0 (Rest) | 0 | 0 | - | - | 1.00 | 0.71 |
| 13 | 8 | 0.026 | 1 | 49 | 0.95 ± 0.05 | 0.50 ± 0.07 |
| 24 | 14 | 0.048 | 1 | 49 | 1.00 | 0.07 ± 0.07 |
| 41 | 24 | 0.082 | 1 | 49 | - | - |
| 59 | 35 | 0.118 | 1 | 49 | - | - |
| 89 | 53 | 0.178 | 1 | 49 | - | - |

**Supplementary Table 2. Parameters used for the simulation of Model 2 and results from the fit.** Parameters in the columns as defined in Extended Data Table 1. Values in column 3-7 calculated according to Model 2 as detailed in the text. For $T_p$ 41, 59 and 89 kPa no [$f_{OFF,C}$; $f_{OFF,D}$] pair exists that allows fitting mean values of all three parameters

| $T_p$ (kPa) | $N_A$ | $f_A$ | $n_i$ | $n_e$ |
|---|---|---|---|---|
| 0 (Rest) | 0 | 0 | - | - |
| 13 | 8 | 0.22 | 44 | 49 |
| 24 | 14 | 0.22 | 39 | 49 |
| 41 | 24 | 0.22 | 32 | 49 |
| 59 | 35 | 0.22 | 24 | 49 |
| 89 | 53 | 0.22 | 11 | 49 |

**Supplementary Table 3. Parameters used for the simulation of Model 3.** Parameters in the columns as defined in Extended Data Table 1. Values in columns 3-5 calculated according to Model 3 as detailed in the text. For all $T_p$ no [$f_{OFF,C}$; $f_{OFF,D}$] pair exists that allows fitting mean values of all three parameters.
